# Spatial point pattern analysis identifies mechanisms shaping the skin parasite landscape in *Leishmania donovani* infection

**DOI:** 10.1101/2021.05.07.443107

**Authors:** Johannes S. P. Doehl, Helen Ashwin, Najmeeyah Brown, Audrey Romano, Samuel Carmichael, Jon W. Pitchford, Paul M. Kaye

**Author notes:** Correspondence to: P.M.K. Vector Molecular Biology Unit, Laboratory of Malaria and Vector Research, National Institute of Allergy and Infectious Diseases, National Institutes of Health, 12735 Twinbrook Parkway, Rockville, MD 20852, USA. BILHI Genetics, 60 Avenue André Roussin, 13016, Marseille, France (Audrey Romano).

## Abstract

Increasing evidence suggests that infectiousness of hosts carrying parasites of the *Leishmania donovani* complex, the causative agents of visceral leishmaniasis, is linked to parasite repositories in the host skin. However, a detailed understanding of the dispersal and dispersion of these obligatory-intracellular parasites and their host phagocytes in the skin is lacking. Using endogenously fluorescent parasites as a proxy, we apply image analysis combined with spatial point pattern models borrowed from ecology to characterize dispersion of parasitized myeloid cells (including Man^R+^ and CD11c^+^ cells) and predict dispersal mechanisms in a previously described immunodeficient model of *L. donovani* infection. Our results suggest that after initial seeding of infection in the skin, heavily parasite-infected myeloid cells are found in patches that resemble innate granulomas. Spread of parasites from these initial patches subsequently occurs through infection of recruited myeloid cells, ultimately leading to self-propagating networks of patch clusters. This combination of imaging and ecological pattern analysis to identify mechanisms driving the skin parasite landscape offers new perspectives on myeloid cell behavior following parasitism by *L. donovani* and may also be applicable to elucidating the behavior of other intracellular tissue-resident pathogens and their host cells.

## Introduction

Vector-borne kinetoplastid parasites of the *Leishmania donovani* complex are the causative agents of zoonotic and anthroponotic visceral leishmaniasis (VL). *Leishmania* parasites occur in two distinct stages; as extracellular promastigotes in their insect vector, hematophagous phlebotomine sand flies (Diptera: Psychodidae: Phlebotominae)^1^, and as intracellular amastigotes in the mammalian host^2^. VL is associated with 3.3 million disability-adjusted life years (DALYs)^3^ and ∼20,000 annual deaths in 56 affected countries^4^. There is no available prophylactic vaccine^5^ and available drugs are either expensive and/or show significant toxicity^6^.

Infection of a naïve host with *Leishmania* parasites occurs through the bite of infected sand flies into the skin, the only interface between insect vector and host. *L. donovani* disseminate from the skin to deeper tissues including the spleen, liver and bone marrow causing systemic pathology, but the skin is commonly asymptomatic^7^. Blood parasite load (parasitemia) is considered the main determinant of outward transmission potential and as a surrogate measure of host infectiousness in VL^8^. Other studies, however, point to the importance of the skin as an additional reservoir of parasites. In a recent study^9^, the infectiousness of *L. donovani*-infected patients in India as measured by xenodiagnoses was found to positively correlate with spleen parasite load as well as blood parasitemia determined by qPCR. Of note, out of 7 infectious patients that remained infectious after drug treatment, 6 had no detectable blood parasitemia^9^. Furthermore, microbiopsy sampling of skin in a VL-endemic region in Ethiopia indicated that parasites can be detected even in asymptomatic individuals, though multiple biopsies from the same patient were often qualitatively and quantitatively different in terms of parasite load^10^. We previously demonstrated in a RAG mouse model of VL that despite high parasite burden in deep tissues, parasitemia was comparatively low (2,000 – 2,500 parasites / ml blood or equivalent to ∼2 – 3 amastigotes / blood meal) and not correlated with outward transmission success^11^. Collectively, these data, together with studies in the hamster model of VL^12^, in canine VL^13^ and in human post-kala azar dermal leishmaniasis^9, 14, 15^ support the need to consider skin parasite load as a potential contributor to infectiousness across the disease spectrum attributable to the *L. donovani* complex.

A confounding feature associated with sampling in many studies relating host infectiousness to skin parasite residence is the possibility of spatial heterogeneity in parasite distribution. Using an immunodeficient model of *L. donovani* infection in mice, we recently demonstrated that obligatory intracellular amastigotes accumulate in patches in asymptomatic host skin, where their dispersion is a strong predictor for the infectiousness of an asymptomatic host^11^. However, a detailed understanding of the mechanisms involved in the micro- and macro-scale dispersion of *Leishmania* parasites (and de facto, of parasitized host cells) in the skin is lacking. As a starting point to deciphering these complex processes, we have further exploited the RAG model of VL, where the absence of acquired immune pressure allows underlying innate pathways controlling the dispersion of parasitized myeloid cells to be more clearly studied.

Here, we describe in detail the micro- and macro-scale dispersion of tandem-dimer Tomato (tdTom) *L. donovani* parasitized cells in the skin of 23 long-term infected RAG mice. Due to the intracellular nature of *Leishmania* amastigotes, their location is equivalent with that of their host cells. Thus, detection of amastigotes can serve as a proxy for detection of parasitized myeloid cells in vivo. We used image analysis tools to extract data based on tdTomato-signal from whole skin images (ImageJ) and also from microscopy images of punch-biopsy skin sections (StrataQuest; see Methods). We then adopted spatial point pattern methodologies from ecology and other fields to interrogate signal dispersion in silico and to make predictions about modes of dispersal^16^. Our results suggest that patches representing infected myeloid cells form “clusters” with a larger patch at the cluster center surrounded by smaller patches. These clusters are consistent with reseeding *L. donovani* amastigotes from a larger patch within a locally varying radius around it. Our data therefore suggest that in the absence of acquired immunity, these patches are the main driver of their own dispersal. This process would also serve to enhance patch growth, as larger patches consist of a merger of smaller patches. This study provides new insights into the complex mechanisms underlying the dispersal and dispersion of *L. donovani*-infected myeloid cells in the skin and exemplifies an approach that may help understand dispersal and dispersion of phylogenetically diverse intracellular pathogens in the skin and other tissues.

## Results

### Parasites accumulate in phagocytic cells in the reticular dermis

We generated confocal images of 20 µm thick longitudinal skin biopsy sections from four different RAG mice (RAG19-22). Visual image inspection suggested that *L. donovani* amastigotes resided primarily in the dermis (**Fig. 1A**). For confirmation, two distinct approaches were used. We calculated the shortest distance measurements of individual amastigotes to the epidermis (**Fig. 1B**) and conducted computationally aided amastigote counting in each skin layer (**Fig. 1C**). The former showed that the majority of parasites resided in the reticular dermis, which comprises the deeper 90% of the dermis^17^. The latter showed that the dermis harbored on average ∼36x and ∼9x more amastigotes / mm^2^ than the epidermis and hypodermis, respectively.

**Figure 1.**
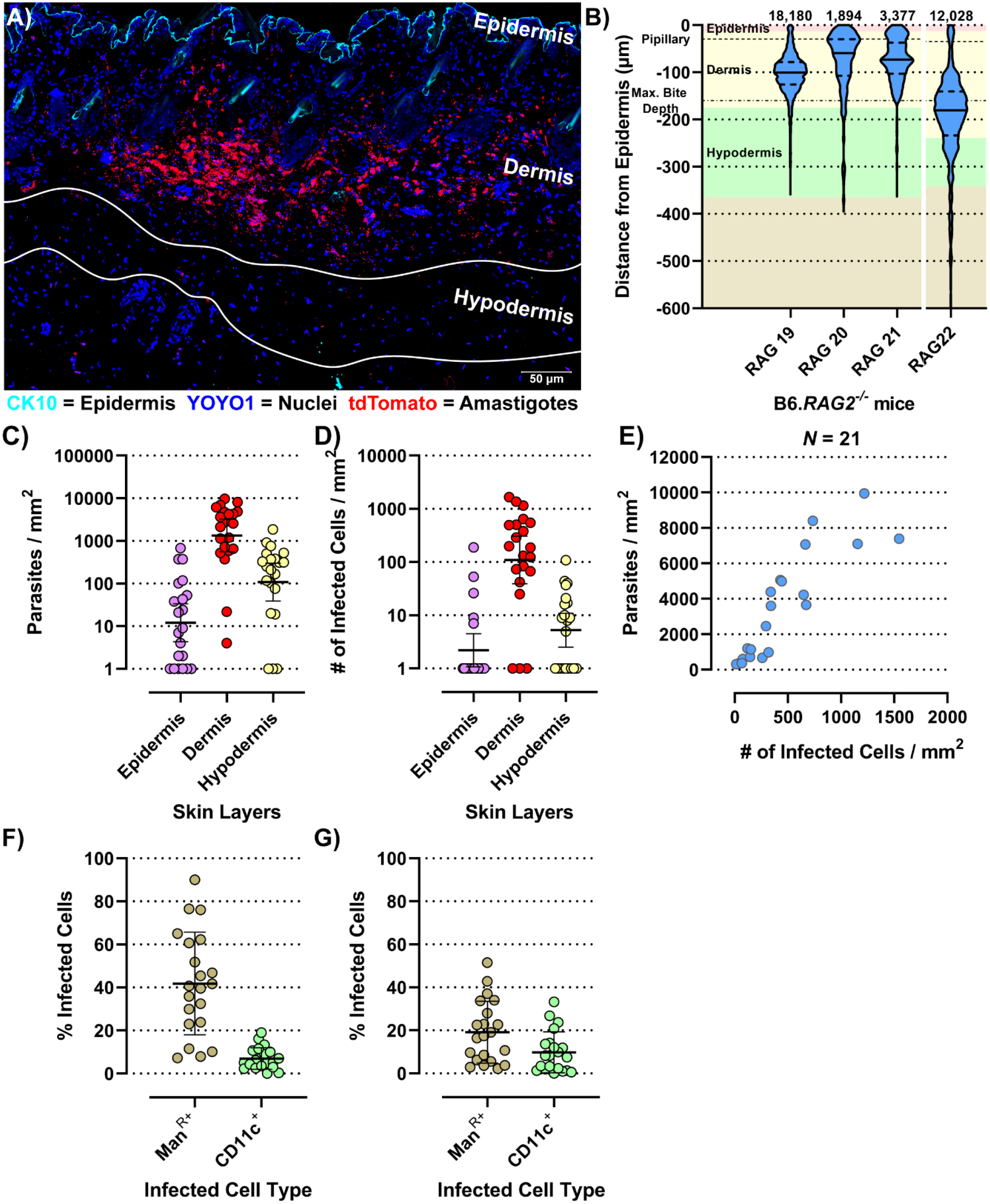
Evaluation of micro-scale parasite distribution in the skin. (A) Representative tile-scan z-stacked confocal microscope image of a 20 µm thick skin biopsy section of the RAG19 mouse. In cyan Cytokeratin-10 (CK-10; epithelium), in blue YOYO-1 (nuclei and kinetoplasts) and in red tdTomato (*L. donovani* amastigotes). The yellow bar is representative of a sand fly proboscis penetrating into the host skin. (B) Violin plot of normalized *L. donovani* amastigote distance from the epithelium. The number of amastigote distances measured are given above each plot. RAG19-21 were female and RAG22 was a male B6.*RAG2^-/-^* mouse. The background color code for skin layer thickness: pink = epidermis, yellow = dermis, green = hypodermis, tan = subcutaneous tissue. The average sand fly proboscis skin penetration depth is marked by dot-dashed line. (C – M) Evaluation of amastigote localization in the skin. Each circle is representative for a measurement from one of 21 analyzed images across RAG19-22. (C) Parasite counts / skin layer (Friedman test: P<0.0001; Dunn’s post hoc test: Epi vs Der: P<0.0001, Epi vs Hypo: P=0.0407, Der vs Hypo: P=0.0047). (D) Counts of infected host cell / skin layer (Friedman test: P<0.0001; Dunn’s post hoc test: Epi vs Der: P<0.0001, Der vs Hypo: P=0.0021). (E) Correlation plot of total parasite counts / skin section against all infected cells / skin section (Spearman r: P<0.0001, r=0.8766, 95% CI: 0.7091 – 0.9505). (F) Percent of detected infected Mannose Receptor positive cells of all infected cells compared to detected infected CD11c^+^ cells of all detected infected cells (Paired t-test: P<0.0001, 95% CI: −33.11 to −18.02). (G) Percent of detected infected Mannose Receptor positive cells of all detected Mannose Receptor cells compared to detected infected CD11c^+^ cells of all detected CD11^+^ cells (Paired t-test: P<0.0001, 95% CI: −11.82 to −4.74).

*Leishmania* amastigotes are obligatorily intracellular parasites, residing in phagocytic cells. Not surprisingly, therefore, infected host cells were also found predominantly in the dermis (∼29x and ∼26x more frequent than in epidermis and hypodermis, respectively; **Fig. 1D**). Thus, parasite dispersion was reflected in host cell dispersion. In fact, the number of parasites / mm^2^ skin correlated very strongly with the number of infected cells / mm^2^ (Spearman r: P<0.0001, r=0.9364, 95% CI: 0.8432 – 0.9749; **Fig. 1E**), suggesting that the proportion of infected cells increased with the increase in parasite burden.

In the skin, macrophages and dendritic cells (DCs) are the most abundant phagocytic cell populations in steady state^18^. Alternatively, activated tissue-resident macrophages (TRM2s) expressing the mannose receptor (Man^R^) are a preferred host for *L. major* in the mammalian skin^19^. CD11c is a generic DC marker^20^, although it can be expressed by tissue macrophages under inflammatory conditions^21^. Thus, Man^R^ and CD11c were used to detect these myeloid cell subsets. We observed that 42% (95% CI: 31% - 52.7%) and 7% (95% CI: 4.7% - 9.3%) of all infected cells were Man^R+^ and CD11c^+^, respectively (**Fig. 1F**). Thus, while Man^R+^ TRM2s also appear to be permissive host cells for *L. donovani* amastigotes, ∼51% of all infected cells detected were neither Man^R+^ or CD11c^+^. Further, of all Man^R+^ and CD11c^+^ cells, 19.1% (95% CI: 12.6% - 25.7%) and 9.8% (95% CI: 5.5% - 14.1%) were infected, respectively (**Fig. 1G**), leaving a large proportion of these cells uninfected (∼80.9% and ∼90.2% of all Man^R+^ and CD11c^+^ cells, respectively). These data suggest that the observed diverse cellular tropism is not due to saturation of permissive targets such as TRM2s.

### Patch landscape is sculpted by parasite reseeding and growth

Whole body skins from 23 untreated, tdTom-*L. donovani* long-term infected (6-7 months) RAG mice (Rag1 – 23) were used to analyze the dispersion of skin parasites within their phagocytic host cell. These RAG mice had been used as control mice in seven independent experiments (Exp 1 – 7) and were comparably treated (**Fig. S1A-C**). We used qPCR-based parasite kDNA detection in multiple skin biopsies from each RAG mouse matched against a standard curve to estimate parasite burden per skin biopsy. Comparing parasite burden from biopsies across the skin of each mouse established whether skin parasite burdens varied geographically across the mouse skin and between mice and the mean parasite burden per biopsy per mouse (**Fig. 2A & S2A**). We found considerable variability in mean parasite burden between RAG mice (Kruskal-Wallis Test: P<0.0001) and in parasite loads between biopsies from the same RAG mouse (**Fig. S2A**) confirming a heterogeneous parasite dispersion in the host skin^11^. Further, data on the parasite burden per biopsy showed in most cases a departure from a normal distribution for each RAG mouse (Shapiro-Wilks test: see **Table S1**) and a lack of equal variance (Brown-Forsythe test: P<0.0001). Thus, the median parasite burden per biopsy was used to estimate the total parasite burden in skin for each RAG mouse. In turn, this was used to estimate the mean parasite burden per cm^2^ of skin (**Table S1**). Despite intravenous inoculation of equivalent numbers of parasites, there was considerable variability in total parasite burden in skin between RAG mice, ranging from ∼1×10^6^ (RAG12) to 1.24×10^9^ (RAG3) parasites per mouse (**Table S1**).

**Figure 2.**
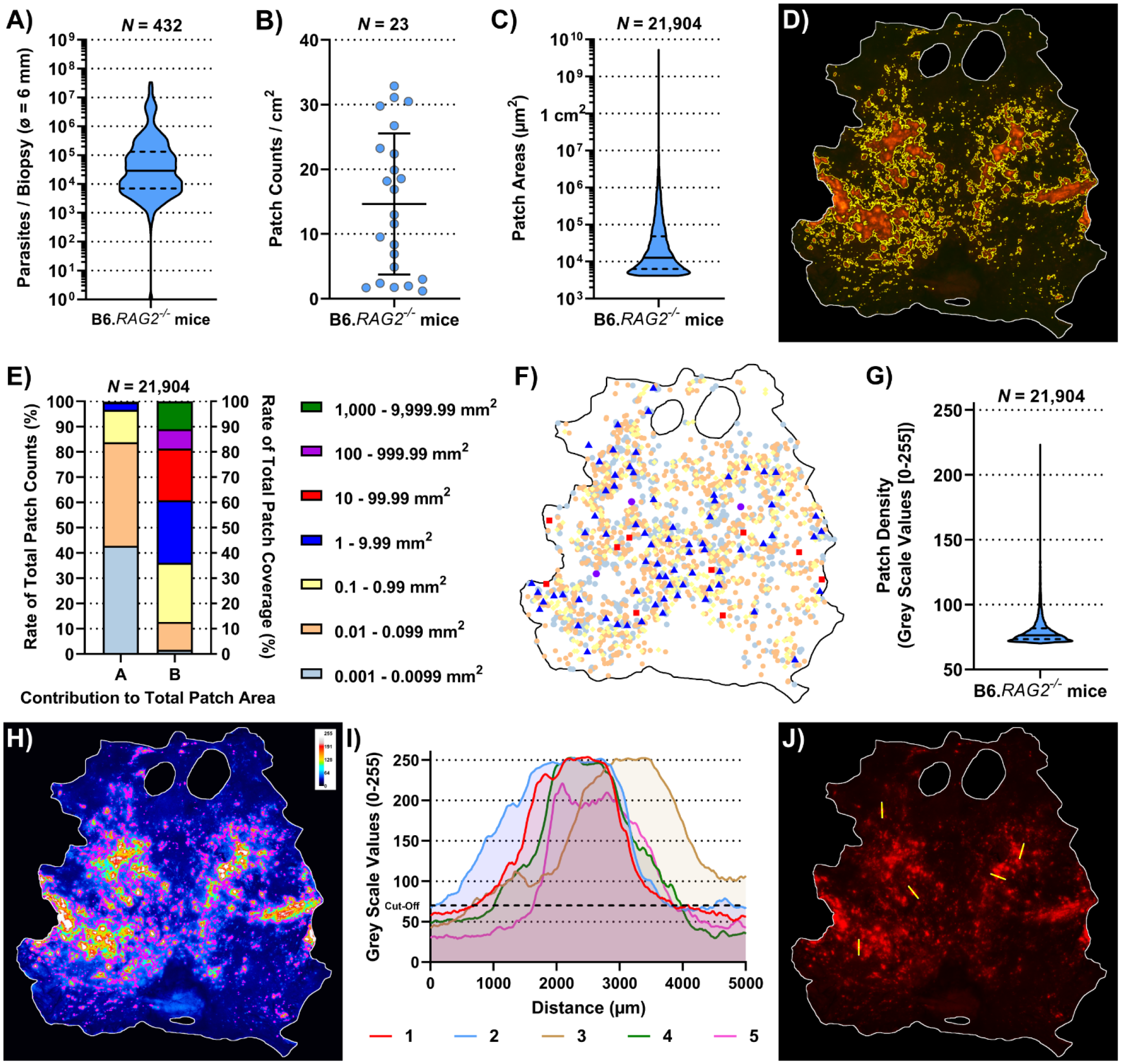
Observational description of the skin parasite landscape. (A) Accumulative qPCR data of estimated parasite loads / biopsy from 432 biopsies (ø 0.6 cm) of 23 RAG mice. (B) Average patch counts / cm^2^ of skin for 23 RAG mice. (I & K) Comparison of ratios of parasitized skin by body region for 23 RAG mice (Friedman Test: P=03113 & P=0.2931, respectively). (C) Accumulative data of the individual area of 21,904 measured patches from 23 RAG mice. (D) Representative image of detected patches in the RAG4 mouse skin. (E) Arranges data from 21,904 individually detected patches from 23 RAG mice arranged by ‘A’ the ratio of patch counts / size category and by ‘B’ the contribution of size category to the overall patched area in the skin. (F) Schematic distribution of patches over the skin of RAG4 marked by patch area according to the categories in (E). (G) Accumulative data of mean patch density / patch of 21,904 individually detected patches from 23 RAG mice. (H) Representative 8-bit grey-scale image showing the tdTomato light intensity per pixel (RAG4). In magenta are the outlines of detected skin patches. (I) X-Y plots of five 5 mm cross sections through patches measuring tdTomato signal intensity / pixel on grey scale [0 – 255] in the RAG4 skin image. The arbitrary cut-off for patch detection set at 70 is marked in these graphs with black dashed line. (J) Image of the RAG4 image indicating where cross sections through patches were made.

Akin to parasite loads, skin patch counts were also highly variable between RAG mice, ranging from ∼1.2 patches per cm^2^ (Rag14) to ∼32.9 patches per cm^2^ (Rag8) and 76 (RAG14) to 2,457 (RAG10) patches per mouse (**Fig. 2B & Table S1**). We also measured the area of skin patches. Patches measured ∼0.00004 cm^2^ (50x smaller than a 25G needle prick) to 53 cm^2^ (>80% of the mouse skin; **Fig. 2C, D, S3 & Table S1**). Most patches were small (<0.1 mm^2^) and only few were large (>1mm^2^), resulting in a heavily skewed dataset (**Fig. S2B**). More precisely, ∼84% detected patches were 0.004 mm^2^ – 0.099 mm^2^ but accounted for merely ∼12.7% of the total patch area (**Fig. 2E**). Conversely, only ∼3.3% of detected patches were >1 mm^2^ but accounted for ∼64% of the total patch area. In an individual mouse, these proportions were even more extreme (**Fig. S4**). In fact, there was a strong correlation between the total patch area and the largest patch in any given mouse (Spearman r: p<0.0001, r=0.9792, 95% CI=0.9496 – 0.9915; **Fig. 3A**). Interestingly, we identified no preferred site of patch accumulation, suggesting that patch dispersion may be random. However, when we plotted patches by size category, we observed that mid- (red and blue symbols; 0.01 – 1 cm^2^) and large-range (violet and green symbols; ≥1 cm^2^) patches were usually surrounded by several small patches (**Fig. 2F & S5**). Moreover, when focusing on mid-range patch dispersion, these patches appeared to follow unidentified tracks across the skin in several RAG mice (**Fig. S5**). These observations suggested that, while patch dispersion may look different between RAG mice, dispersal is not a random process, but governed by unknown factors in the skin. Our subsequent analysis aimed to test this assertion.

**Figure 3.**
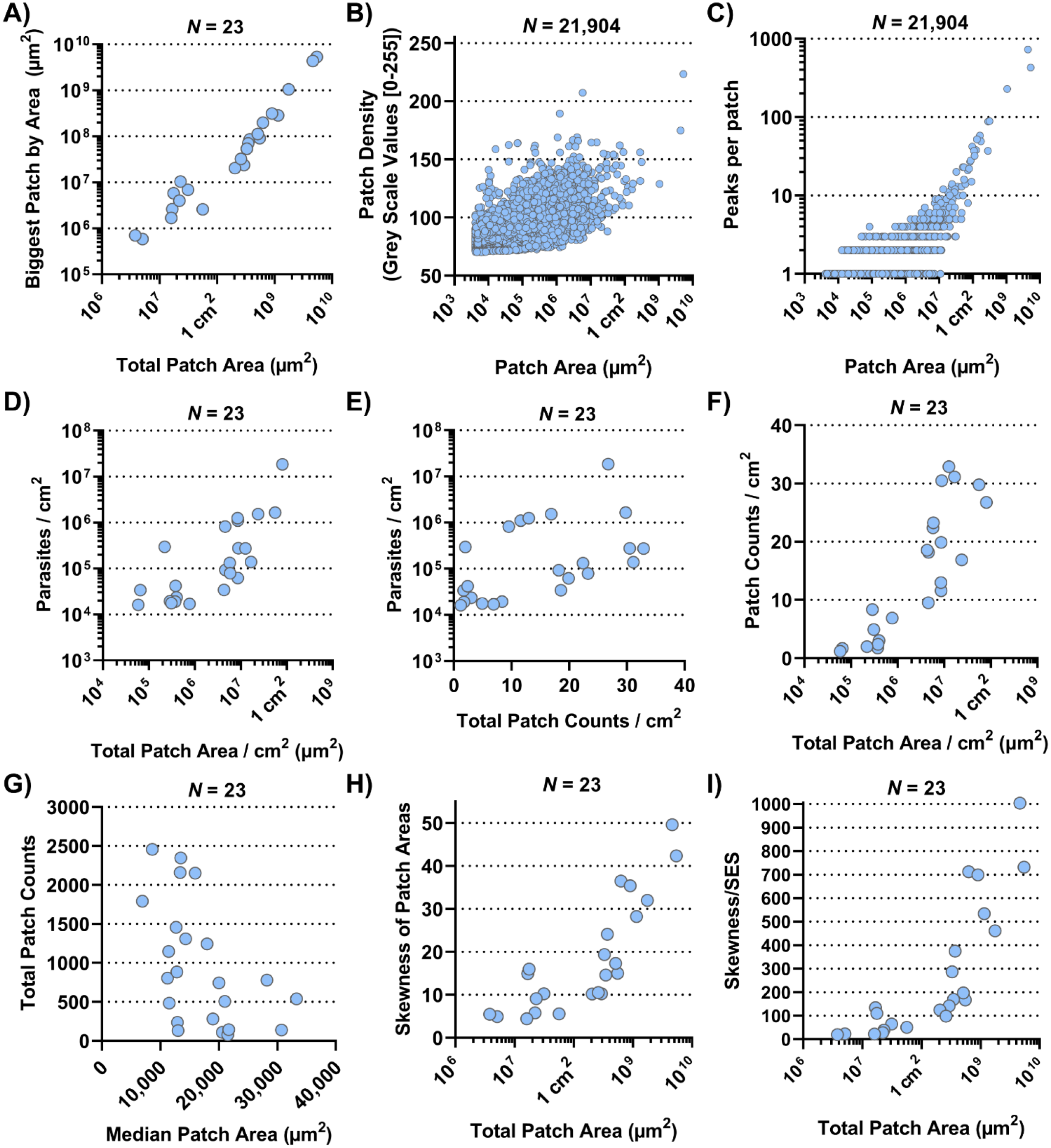
Correlation plots. (A) Correlation plot of total parasite patched skin by area (µm^2^) against the size of the biggest parasite patch detected in each of the 23 RAG mice (both on a log-scale; Spearman r: p<0.0001, r=0.9792, 95% CI=0.9496 – 0.9915). (B) Correlation plot of 21,904 skin parasite patches by area (µm^2^) from 23 RAG mice (on a log-scale) against their respective mean pixel density on a 8-bit grayscale (Spearman r: P<0.0001, r=0.6554, 95% CI=0.6476 – 0.6631). (C) Correlation plot of 21,904 skin parasite patches by area (µm^2^) from 23 RAG mice against the number of high-density peaks per individual patch (both on a log-scale; Spearman r: P<0.0001, r=0.3831, 95% CI=0.3714 – 0.3947). (D) Correlation plot of total parasite patched skin by area (µm^2^) normalized to 1 cm^2^ of skin against parasite burden / cm^2^ for each of the 23 RAG mice (both on a log-scale; Spearman r: P P<0.0001, r=0.7589, 95% CI=0.4948 – 0.8947). (E) Correlation plot of total skin parasite patch counts normalized to 1 cm^2^ of skin against mean parasite burden per punch biopsy for each of the 23 RAG mice (on a log-scale; Spearman r P=0.0021, r=0.5741, 95% CI=0.1997 – 0.8022). (F) Correlation plot of total parasite patched skin by area (µm^2^) normalized to 1 cm^2^ of skin (on a log-scale) against total skin parasite patch counts normalized to 1 cm^2^ of skin for each of the 23 RAG mice (Spearman r: P<0.0001, r=0.8656, 95% CI: 0.6984 – 0.9432). (G) Correlation plot of the median patch area (µm^2^) against total skin parasite patch counts for each of the 23 RAG mice (Spearman r: P=0.006, r=- 0.5148, 95% CI=-0.7701 to −0.1175). (H) Correlation plot of total parasite patched skin by area (µm^2^; on a log-scale) against the skewness of parasite patch area data for each of the 23 RAG mice (Spearman r: P<0.0001, r=0.8686, 95% CI=0.7044 – 0.9445). (I) Correlation plot of total parasite patched skin by area (µm^2^; on a log-scale) against the test statistic that infers skewness to the population for each of the 23 RAG mice (Spearman r: P<0.0001, r=0.9279, 95% CI=0.8311 – 0.9701).

We observed that patches had higher tdTomato signal at their center compared to their periphery. Since tdTomato amastigotes are approximately of equal brightness, it is reasonable to associate the brighter signal at the patch center with higher amastigote density. A grey-scale pixel analysis in the red channel to measure tdTomato signal intensity per pixel (**Fig. 2H & S6**) showed that most mean grey-scale values per patch were close to the cutoff of 70 (**Fig. 2G, Fig. S2C**). However, when patch area was correlated with their respective average grey-scale value, it showed a strong correlation (**Fig. 3B**; Spearman r: P<0.0001, r=0.6554, 95% CI=0.6476 – 0.6631). This suggested that as patches grew outward, they became more densely populated with amastigotes. Cross section analysis of tdTomato signal intensity in selected patches showed volcano-like graphs with steep shoulders, suggesting that the amastigote bulk is concentrated at the center of a patch and then thins out quickly at the patch periphery (**Fig. 2I & J**).

We also observed that larger patches contained multiple areas of high tdTomato-signal (**Fig. S6**). Correlating patch area and density peaks per patch suggested that all mid-range and larger patches had multiple density peaks (Spearman r: P<0.0001, r=0.3831, 95% CI=0.3714 – 0.3947; **Fig. 3C**). The statistically weak correlation was explained by the broad base of small patches with only one or two peaks. Accumulation of density peaks per patch only became apparent when patch areas exceeded 1 mm^2^ (>10^6^ µm^2^). This suggested that smaller expanding patches grew into one another to form bigger patches. Furthermore, large patches indicated areas where initial parasite seeding had been higher and / or earlier.

Further, we investigated the relationships between mean parasite burden / cm^2^, patch counts / cm^2^ and total patch area / cm^2^ by correlation plot analysis. Mean parasite burden / cm^2^ correlated strongly with total patch area / cm^2^ of skin, suggesting that increases in total skin parasite burden would also increase total patch area and thus may be the driver of patch growth (**Fig. 3D**; Spearman r: P<0.0001, r=0.7589, 95% CI=0.4948 – 0.8947). Patch counts / cm^2^ correlated only moderately with parasite burden / biopsy (**Fig. 3E**; Spearman r: P=0.0021, r=0.5741, 95% CI=0.1997 – 0.8022), suggesting that increases in skin parasite burden was more closely related to patch expansion than new patch seeding. Conversely, when total patch area / cm^2^ was correlated with the patch count / cm^2^, it showed a very strong positive correlation (**Fig. 3F**; Spearman r: P<0.0001, r=0.8656, 95% CI: 0.6984 – 0.9432), suggesting that the proportion of parasitized host skin is largely determined by the number of observed patches. Correlating the median of patch areas per skin for each RAG mouse to the total patch counts per skin, we found a moderately negative correlation (**Fig. 3G**; Spearman r: P=0.006, r=-0.5148, 95% CI=-0.7701 to −0.1175), which suggested that the more patches a RAG mouse skin had the more of these patches were likely to be small (<0.1 mm^2^).

While data skewness is an underlying characteristic of skin parasite dispersion for any given sample, skewness cannot be directly inferred onto a population. To test whether the observed sample skewness was a general population feature, we divided sample skewness by the standard error of skewness (SES)^22^. The resulting test statistic being outside the interval [-2, 2] infers that data skewness was a population property (**Table S1**). For instance, total patch area per skin was very strongly correlated with the degree of skewness in the patch area data (**Fig. 3H**; Spearman r: P<0.0001, r=0.8686, 95% CI=0.7044 – 0.9445), suggesting that as patchiness increased in the skin, mid-range patches give way to a few large patches surrounded by many small patches. This appears to be a common population feature, given the correlation of the test statistic to the total patch area / skin was even stronger than compared to the sample skewness (**Fig. 3I**; Spearman r: P<0.0001, r=0.9279, 95% CI=0.8311 – 0.9701).

### The skin patch landscape is heterogeneously clustered

We previously established that the skin parasite dispersion pertained to concepts of landscape epidemiology^11^. Thus, we applied methodologies of spatial point pattern analysis commonly used in landscape epidemiology to characterize further the dispersion of parasites and their myeloid host cells in skin and identify potential modes of their dispersal^16^. We converted our patch data into spatial point patterns by mapping the center of mass of each patch to a window characterized by the respective RAG skin outline (**Fig. 4A, Fig. S7, Table S2**). After verifying the absence of point duplications, characteristics of patches (e.g. area and density) could optionally be reintroduced to the pattern as a mark (**Fig. 4B, Fig. S8**). Focusing on patch area, we calculated the intensity of point distributions (equivalent to dispersion) with or without a mark and graphically represented the measured point pattern intensities (**Fig. 4C, D, S9 & S10**). Alternatively, we calculated the intensity per cm^2^, which is akin to the number of patches per cm^2^ and found that both methods corroborated one another (**Table S1 & S2**). As shown in **Figure 4C** (without area mark), intensity showed the distribution of patches by center of mass location, whereas in **Figure 4D** (with area mark) intensity showed the distribution of patches by size in the skin (see also **Fig. S9 & S10**). Both intensities showed a clear departure from a homogeneous distribution. To gain further understanding of how patches related to one another, we performed a cluster analysis, choosing 3.9 mm as the maximum distance between patch borders (see Methods). This showed small patches clustering strongly around larger patches (**Fig. 4E & S11**), confirming our previous observation from **Figure 2F** (see also **Fig. S5**). Further, this analysis showed the formation of networks of closely clustered patches, which may enhance the likelihood of parasite pick up by a biting sand fly without having to densely parasitize the whole skin (**Fig. 4E & S11**).

**Figure 4.**
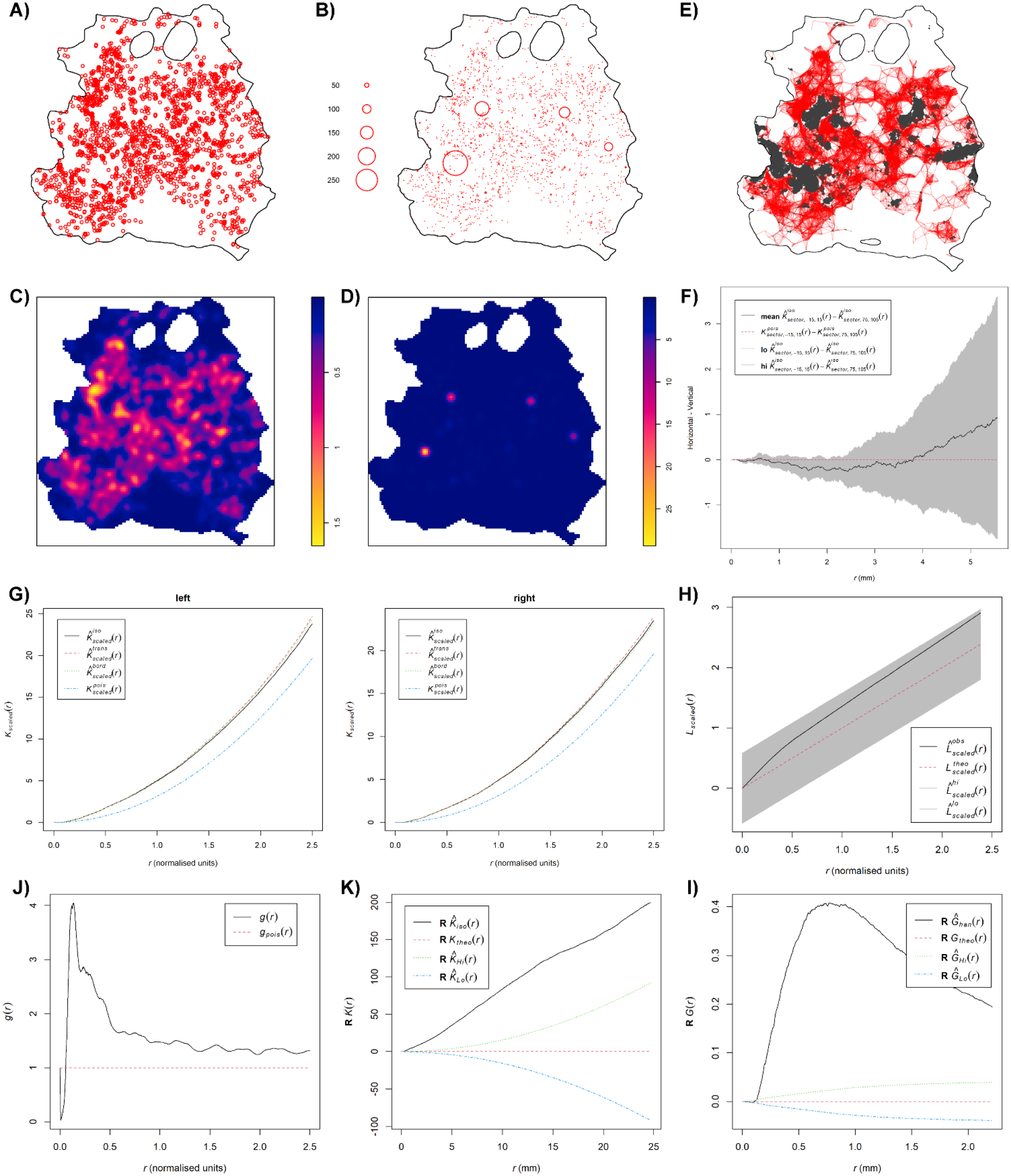
Spatial point pattern analysis of skin parasite patch distribution. All graphs are examples for spatial point pattern analysis based on the RAG4 data. (A) Spatial point pattern of skin parasite patches based on center of mass of each detected patch without respective marks. (B) Spatial point pattern of skin parasite patches based on center of mass of each detected patch with respective patch area mark. (C) Patch intensity representation of skin parasite patches based on center of mass of each detected patch without respective marks. (D) Patch intensity representation of skin parasite patches based on center of mass of each detected patch with respective patch area mark. (E) Representation of a cluster analysis output linking all patches whose patch borders are ≤3.9 mm into a network. (F) Graphical representation of the isotropy test showing the theoretical Poisson process (red dotted line), the RAG4 data (black solid line) and the global envelopes (grey). (G) Graphical representation of correlation-stationary test based on a locally scaled K-function. (H) Scaled L-function showing the theoretical Poisson process (red dotted line), the RAG4 data (black solid line) and the global envelopes (grey). (I) Pair correlation function showing the theoretical Poisson process (red dotted line), the RAG4 data (black solid line). (K) Data points independence test based on K-function of the data residuals, showing the theoretical Poisson process (red dotted line), the RAG4 data (black solid line) and the upper (green) and lower limits (blue) of the confidence intervals of the theoretical Poisson process.

These observations suggested that patch dispersion is likely to be neither homogeneous nor random. To formally test for homogeneity of patch distributions, a quadrat test based on the Fisher’s exact test was applied, which rejected homogeneity for all patch distributions (P=0.001; **Table S2**). Thus, an inhomogeneous Poisson point process (IPP) was used as a null hypothesis. Analogue to a homogeneous Poisson point process, an IPP is assumed to be correlation-stationary (the pair correlation between two spots in the window depends only on their relative positions) and isotropic (the distribution is unaffected by the pattern orientation). However, isotropism was rejected for about half of all patch patterns (**Fig. 4F & S12**); RAG14 did not produce a reliable test result due to its sparse number of patches. To assess correlation-stationarity, we split the point patterns into two portions of equal numbers of observations and applied the inhomogeneous K-function, which is derived from Ripley’s K-function, to each half of the pattern. The analysis showed considerable disagreement between the two plots for several mice indicating that the whole point pattern was not correlation-stationary (**Fig. S13**). In addition, the inhomogeneous K-function showed considerable disagreement between different applied border corrections, which suggested that an IPP was a bad fit to describe patch patterns. In addition, we rejected a Poisson distribution for our patch patterns based on a Quadrat Count test (**Table S2**). These disagreements suggested that the spatial scale of point interactions, which is assumed to be constant by the inhomogeneous K-function^16^, was locally variable for our patch patterns. Thus, by assuming a point process that was locally stationary and isotropic but showed a varying rescaling factor between locations in the skin, we could use a locally scaled K-function to rerun the correlation-stationary test. Although not perfect, agreement between the two plots per mouse and between different border corrections was significantly improved (**Fig. 4G & S14**), suggesting that patch distribution was locally scaled.

We proceeded with the scaled L-function, derived from Besag’s L-function, to determine if patches were evenly spaced (regular), randomly spaced (independent) or clustered (**Fig. 4H & S15**). While the fitted data (black solid line) generally departed from the hypothetical random distribution toward a clustered one (black solid line above dotted red line), only RAG11, RAG13, RAG14 and RAG23 showed a clear departure from the global simulation envelops of a hypothetical random distribution (grey shaded area) toward clustering. To confirm these results, we used the pair correlation function (PCF). The scaled PCF analysis showed for most mice, patch patterns departed from the theoretical random distribution (dotted red line) toward a clustered point process (black line above the red dotted line), in particular, at short distances between patches (**Fig. 4J & S16**). With increased distances between patches, some point patterns showed an approximation to more point independence. Applying the K- and nearest-neighbor (G-) function to the point pattern residuals showed a clear departure of independence toward clustering at shorter distances, indicating positive point dependence (black line outside the upper confidence intervals [dotted green line]) for both tests (**Fig. 4K, L, S17 & S18**).

### Modelling the point process behind patch patterns suggests influence of covariates

Consequently, to model patch distribution we adopted a model that accounted for positive point (patch) dependence (clustering) and was tolerant to a locally varying distribution scale. We used different cluster processes (Cauchy, Matérn, Thomas and Variance-Gamma) as described by a Baddeley et al. (‘spatstat’^16^) that assume that cluster formation is based on a parental point distribution that seeded the observed points around themselves within a radius, which is variable between clusters due to the locally scaled nature of cluster distribution. This assumption is reflected in our data showing that small patches cluster around mid-range and large patches (**Fig. 2F, 4E, S5 & S11**).

13 of the 23 patch distributions were best fitted by a Matérn cluster process, while the other 10 were best fitted by a Thomas cluster process. The Matérn cluster process assumes a uniform distribution of offspring around the parental point, while the Thomas cluster process assumes offspring to be randomly displaced from the parental point^16^. Checking the parameters selected by the model showed that they were within a reasonable range, confirming model validity (**Table S3**). This was further confirmed by the scaled L-function of the selected fitted models for each patch pattern (**Fig. 5A & S19**). The L-functions of the fitted models closely resembled the L-functions from the point pattern analyses (**Fig. 4H & S15**). Further, we used the selected fitted models to run 4 simulations each to see if the outcome would resemble the distributions the original data (**Fig. 5B & S20**). With a few exceptions, like RAG3, the simulation outputs provided a good representation of the original patch dispersion data (**Fig. 4A & S7**).

**Figure 5.**
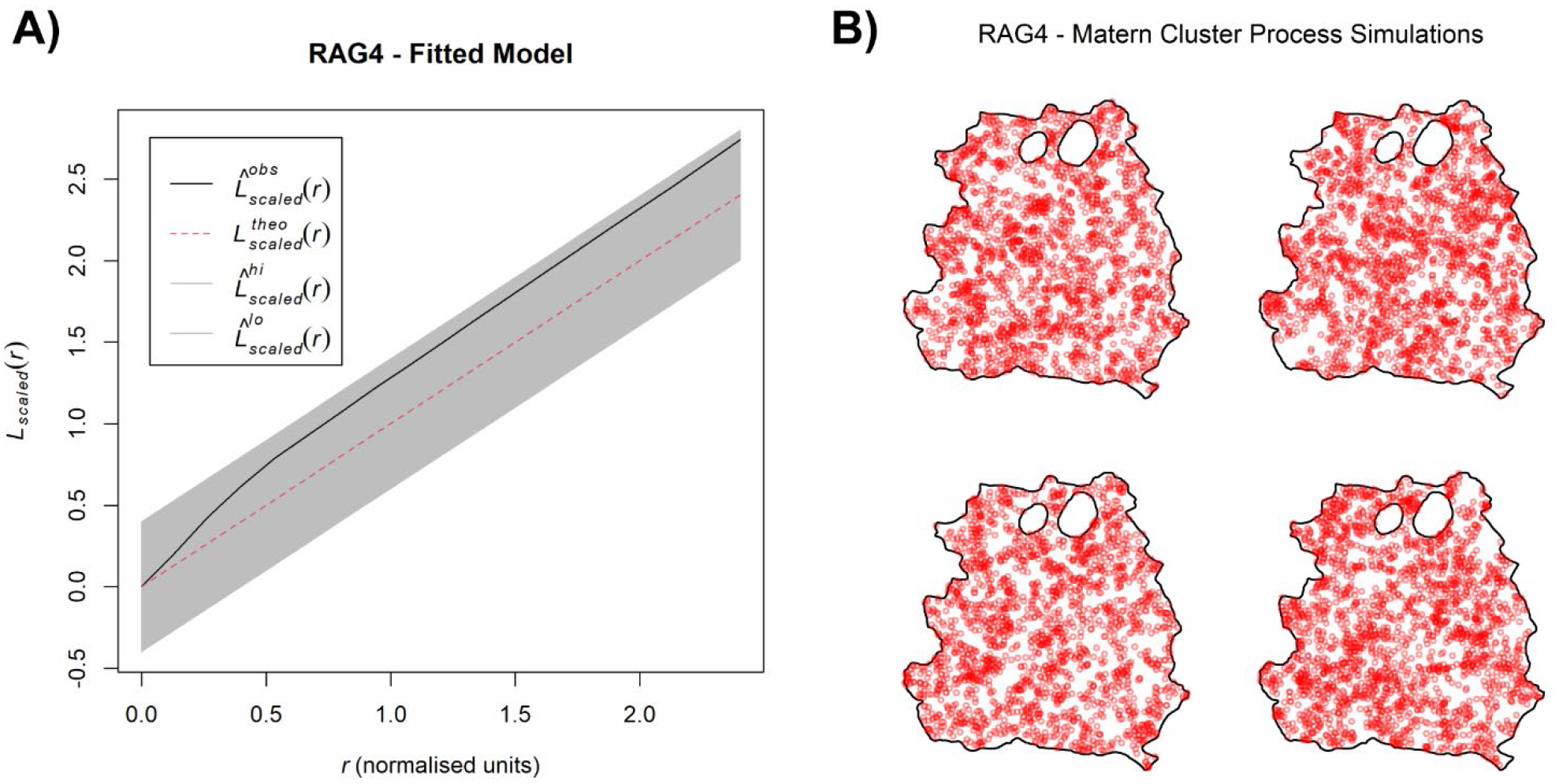
Basic model evaluation of skin parasite patch distribution. (A) Scaled L-function showing the theoretical Poisson process (red dotted line), the RAG4 predicted model data (black solid line) and the global envelopes (grey). (B) Four model simulations based on the Matérn cluster process for RAG4.

Based on these findings, we can deduce how the dispersal of parasites and their myeloid host cells may operate. Both the Matérn and Thomas cluster processes assume points of origin that seed new points within a set radius to create an observed clustered point dispersion. In a locally scaled setting, as it is the case for our patch distribution, this radius is locally varying in length and thus distinct for each patch cluster. Analogously, after initial skin infection, the dispersal of patches may be explained by existing patches locally seeding new patches around themselves, while growing outward, creating a self-propagating network of patch clusters in the skin (**Fig. 6 & S21**).

**Figure 6.**
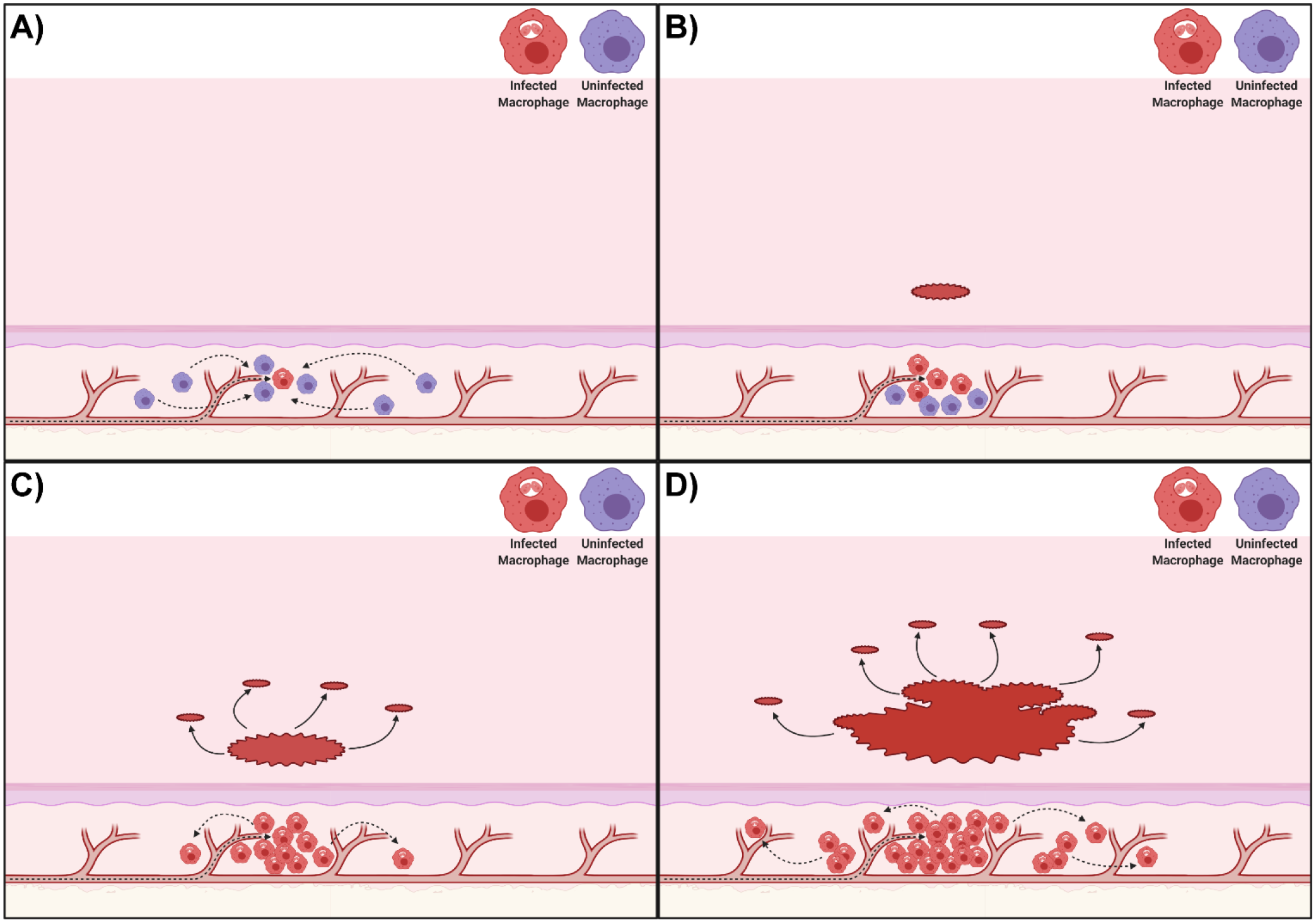
Proposed mechanism for the dispersal of patches of *L. donovani*-infected myeloid cells. (A) After an initial *Leishmania* amastigote seeding event in the skin e.g. by arrival of an infected myeloid cells (in red) from the circulation, uninfected phagocytic cells (in purple) in the skin are attracted to the infected cells to form a type of innate granuloma. (B) Uninfected myeloid cells are gradually infected in the innate granuloma forming a patch of infected myeloid cells. (C) As these patches keep growing, they seed new patches within a radius ‘r’ around themselves, potentially by escape of infected myeloid cells or via transient release of free amastigotes. (D) This process then keeps repeating itself, forming self-propagating networks of patch “clusters”. All images were created with BioRender.com.

## Discussion

It is known that heterogeneity in skin parasite dispersion provides a mechanism to enhance host infectiousness to sand fly vectors^11^. Here we have provided evidence for the underlying mechanisms driving this heterogeneity. To our knowledge, this is the first study to make use of spatial point pattern methodologies^16^ in this way. The Matérn and Thomas cluster processes, which were identified to best describe how dispersion of patches of parasites and their infected myeloid host cells occurred, both assume points of origin which seed new points within locally varying radii clustering around them. Based on this analysis, we therefore propose that existing patches locally seed new patches around themselves, giving rise to patch clusters with a central large patch. Since patches are densely parasitized, these patch networks would increase the chance of high parasite loads being acquired by a sand fly and eliminate the need for parasitism of the entire skin. Indeed, our conservatively chosen tdTomato-signal and patch size thresholds may underestimate very small patches leading to patch networks that are denser than reported here, further enhancing outward transmission potential.

In vivo, *Leishmania* amastigotes reside in phagocytic cells in the dermis, primarily macrophages^23^. Therefore, parasite dispersion is linked to host cell dispersion, a subject on which scant information is available. In the steady-state dermis, macrophages and DCs are the primary phagocytic cell populations^18^. Interestingly, macrophages and DCs have very particular distributions in steady-state skin. Branched DCs are predominantly interstitially localized directly underneath the dermal-epidermal junction^24, 25^. Macrophages are commonly found below these DCs; mostly interstitially. Deeper in the reticular dermis, both cell types associate perivascular to blood vessels rather than lymphatic vessels and DCs here are more rounded. Among the different macrophage populations in the skin, it was shown that Man^R+^ TRM2s are important as long-term *L. major* host cells^19^. In addition, the rate of parasite infection in Man^R+^ TRM2s (∼45%) correlated strongly with the severity of skin pathology and parasite survival in the host. The rate of infection in DCs was much lower (22.5%), indicating a preference of *L. major* for Man^R+^ TRM2s^19^. In our study, the observed infection rates for both cell types with *L. donovani* were less than half of this (∼19.1% and 9.8%, respectively), but *L. donovani* similarly showed a preference for Man^R+^ TRM2s over DCs (5:1). These cells constituted almost 50% of all infected host cells. Conversely, 50% of host cells belonged to neither group; presumably, other myeloid-derived phagocyte populations^23^. Interestingly, the majority of Man^R+^ TRM2s and CD11c^+^ DCs (∼80.9% and ∼90.2%, respectively) were uninfected in the analyzed skin sections, despite the high number of parasites present in some of the sections. This raised the question of why the small proportion of infected host cells should be clustered close together and also indicates that observed cellular tropism is not due to saturation of preferred targets.

Further, our analysis suggested that the process of forming patches may be influenced by yet-to-be-identified covariates. One covariate may be proximity to blood vessels as Man^R+^ TRM2s often colocalize to them in the dermis, where they can pick up macromolecules from the blood^19, 26^. Given that parasitized phagocytic cells in circulation may have properties associated with apoptotic cells^27^, it is possible that TRMs capture parasitized host cells by efferocytosis, thus seeding new skin sites for patch formation. Alternatively, infected phagocytes in circulation could exit the vasculature by extravasation at sites of sub-clinical inflammation. Either mechanism could explain why mid-range patch distribution appeared to follow unidentified lines in the skin and why there was no preferred location for parasite accumulation in the skin. Further, proximity to blood vessels may also enhance access to nutrients from the circulation for parasitized phagocytic cells and thus, indirectly promote parasite proliferation. Also, parasite proximity to blood vessels would improve their chances to be found by a probing sand fly. Future analysis could potentially confirm blood vessel association of parasite patches by proximity measurements. Three-dimensional (e.g. light sheet) microscopic imaging data would be needed to achieve this. This and other covariates that may affect the establishment of parasite niches can be integrated in the future into the Matérn and Thomas cluster process models to determine their influence on the dispersal and dispersion of parasites and their myeloid host cells.

Independent of the means of initial skin seeding of parasites, our data indicate that the accumulation of infected myeloid cells into patches could be analogous mechanistically to phagocyte migration during granulomatous inflammation. For example, early in the process of *L. donovani*-dependent granuloma formation in the liver, uninfected Kupffer cells (KCs) migrate towards a granuloma-initiating infected KC resulting in a heterogeneous distribution of KCs and their condensation and subsequent infection at the granuloma core^28, 29^. Conversely, studies in zebrafish embryos infected with *Mycobacterium marinum* highlight the innate capacity of phagocytes to exit granulomas^30^, an innate feature of the phagocytic cell response to infection that appears to be obscured in fully immunocompetent mouse models of granulomatous inflammation^28^. Our RAG mouse model of skin infection similarly lacks the confounding effects of an adaptive immune response, providing a similarly unique window on processes driving innate myeloid cell aggregation and a platform on which to build increasing levels of immune complexity. It remains to be determined to what extent patches form because of the inherent migratory properties of phagocytic cells and / or modification of such properties by intracellular amastigotes. Manipulation of phagocyte function by intracellular *Leishmania* has been well-described^31^ though few studies have addressed phagocyte migratory potential. Similarly, whilst phagocyte heterogeneity in skin is well described (see above), the mechanisms regulating mobility of these cells is their natural 3D-environment remain poorly understood.

The notion that new patches are seeded primarily from existing patches is supported by the observation that larger patches were surrounded by smaller patches that emerged within limited distances from the larger patches, resulting in patch clustering. If, akin to parasite dispersion, parasite dispersal is also linked to its host cells, then, mechanistically, patch clustering could occur by escape of infected host cells from these “innate granuloma”-like parasite patches as was demonstrated in *M. marinum*-infected zebrafish embryos^32^. Conversely, parasites could be redistributed to uninfected host cells within a limited radius from patches by amastigote release from ruptured host cells or by intercellular passage. Macrophages are known to form a variety of open-ended tunneling nanotubes (TNT) between one another at a distance and such structures have been implicated in the spread of several respiratory viruses and HIV-1^33–35^. We know of no formal evidence, however, to show that *Leishmania* parasites are transported between macrophages via TNTs.

The cores of patches are very dense with parasites and heavily infected host cells, but the patch fringes are much less heavily parasitized, suggesting that parasites proliferating strongly within a patch are then pushed outward. Mandell & Beverley (2017) observed that not all amastigotes in the host skin proliferate at the same rate and described two distinct amastigote populations; one that was dormant, the other proliferative^36^. Thus, patches may represent areas of where amastigotes are proliferative, while inter-patch space could mark areas of dormant amastigotes. Whether host cell metabolism stimulates/supports amastigotes to proliferate in the mammalian host skin^37^, remains to be determined.

In summary, through spatial point pattern analysis combined with systematic image analysis we provide a novel mechanistic framework to explain how the dispersal of *L. donovani* infected myeloid cells occurs in the absence of the constraints imposed by acquired immunity.

## Materials and Methods

### Ethics statement

All animal usage was approved by the University of York Animal Welfare and Ethics Review Committee and the Ethics Review Committee at FERA Science Ltd., and performed under UK Home Office license (‘Immunity and Immunopathology of Leishmaniasis’ Ref # PPL 60/4377).

### Mouse, Leishmania and sand fly lines

C57BL/6 CD45.1.*Rag2^-/-^* (RAG) mice (originally obtained from the National Institute of Medical Research, London, UK) were used. All animals were bred and maintained at the University of York according to UK Home Office guidelines. The Ethiopian *Leishmania (Leishmania) donovani* strain (MHOM/ET/1967/HU3) had been transfected with a tdTom gene^38^ to generate a tdTom-*L. donovani* line in another study^29^. The parasites were maintained RAG mice by repeat-passage of amastigotes. For both parasite passage and RAG mouse infection, tdTom-*L. donovani* amastigotes were isolated from the spleens of long-term infected RAG mice and i.v. at 8×10^7^ amastigotes / uninfected RAG mouse as previously described^11^.

### Mouse skin harvest

All animals were untreated control animals pooled from 7 different experiments. All animals were exposed to comparable experimental conditions (**Fig. S1A & B**). Mouse skin harvest was performed as previously described^11^. Briefly, RAG mice were sacrificed in a CO_2_-chamber, followed by cervical dislocation. Animals were first shaven with electric clippers and then treated with a depilation cream (Veet by Reckitt, Slough, UK) for up to 3 min. The depilation cream was scraped off and residual cream was washed off under running water. Mice were skinned by an abdominal double-Y incision, removal of ears and exclusion of paws and tail (**Fig. S1C**). The skin was pulled from the rump over the head and then cut off at eye-level. Excess adipose tissue was removed with curved tweezers and skins were kept in complete RPMI on ice until stereomicroscopic imaging and subsequent biopsy punching.

### Genomic DNA extraction

16 or 24 skin biopsies (ø 0.6 cm) were taken from each skin as previously described^11^. All genomic DNA (gDNA) extractions were performed with either the DNeasy^®^ Blood & Tissue spinning column kit (Qiagen, Venlo, Netherlands) or the DNeasy^®^ 96 Blood & Tissue plate kit (Qiagen) according to supplier’s protocol.

### Quantitative Polymerase Chain Reaction

All quantitative real-time PCRs (qPCR) were performed as previously described^11^. Briefly, previously characterized *Leishmania*-specific kinetoplastid DNA primers (Accession number AF103738)^39^ were used at a final concentration of 200 nM. 2 ng of total gDNA extracted from skin biopsies were used / reaction. Fast SYBR^®^ Green Master Mix (Applied Biosystems) was used according to supplier’s guidelines. Reactions were run in a StepOnePlus™ thermal cycler (Applied Biosystems, Waltham, MA, USA) with a thermal cycle of 95°C for 20 s, a cycling stage of 40 cycles of 95°C for 3 s, 60°C for 5 s, 72°C for 20 s, 76.5°C for 10 s (data read at final step), followed by the standard melt curve stage. Data was analyzed by StepOne™ Software v.2.3.

### Microscopy

Fluorescent stereomicroscopy of whole skins was performed as previously described^11^. Briefly, a series of 30-40 images at 12x magnification were taken / skin to image the whole skin area from the hypodermal face. All images were taken under the same conditions (exposure time: 400 ms). Image-series were collated in Adobe^®^ Photoshop^®^ to render a single image of the whole mouse skin (Abode Inc., San Jose, CA, USA).

For confocal microscopy, fixation and cryo-preservation of skin punch biopsies (ø 0.6 cm) was adapted from Accart et al. (2014) to optimize tdTomato-signal and skin integrity preservation after skin freezing and cutting^40^. Skin biopsies were fixed in 2% Formaldehyde in PBS for 15 min at room temperature (RT), rinsed 3x in 15% Sucrose/PBS and incubated overnight at 4°C in a Tris-Zinc fixative (per 100 ml: 1,211.4 mg Tris-base, 501.5 mg ZnCl_2_, 500.5 mg Zinc Acetate Dihydrate, 50.6 mg CaCL_2_, 15 g Sucrose dissolved in dH_2_O) on a shaker. The next day, biopsies were rinsed 3x in 15% Sucrose/PBS and incubated in 7.5% Porcine Gelatine (300 bloom) in 15% Sucrose/PBS in individual cryo-molds at 37°C for 1 h. Biopsies were sealed upright in the gelatine blocks by congealing at RT. Gelatine/specimen-blocks were cut down in size, flash-frozen in isopentane on dry ice, further frozen in OCT in a cryo-mold on dry ice and stored at −80°C. Frozen skin biopsies were longitudinally cut at 20 µm thickness to expose all skin layers and section were placed on superfrosted microscopy glass slides. Gelatine was removed by slide incubation in PBS at 37°C for up to 30 min.

Antibody-labelling of skin sections was adapted from Dalton et al. (2010)^41^. Briefly, slides were rinsed in PBS, permeabilized for 15 min with 0.3% Triton X-100/PBS (perm. buffer), rinsed in PBS, blocked for 30 min with Image-iT™ FX Signal-enhancer (Invitrogen, Waltham, MA, USA), rinsed with PBS and further blocked for 30 min with 5% serum + FcBlock (1:1000) in perm. buffer. Primary antibodies were diluted in 5% BSA in perm. buffer (CK-10 [1 µg/ml] (Abcam: ab76318), Mannose Receptor [1 µg/ml] (Abcam: ab64693), CD11c [5 µg/ml] (Abcam: ab33483)) and incubated with specimen for 90-120 min at RT (Abcam PLC, Cambridge, UK). Slides were washed 3 times in 0.5% BSA/PBS. Secondary antibodies were diluted in 5% BSA in perm. buffer (BV421 [1 µg/ml] [Biolegend: 406410]; Alexa Fluor 647 [Invitrogen: A-21451]) and incubated for at least 60 min at RT (BioLegend, San Diego, CA, USA). Slides were washed 2 times in 0.5% BSA/PBS and once in PBS at RT. Specimen were counterstained with 0.2 µM YOYO-1 in PBS for 30 min at RT and washed 2 times in PBS. Slides were mounted in ProLong Gold (ThermoFisher Scientific, Waltham, MA, USA) and left overnight at 4°C. Slides were sealed the next day and imaged with a 40x oil-immersion objective (400x magnification) on a fully motorized invert LSM 880 confocal microscope with Airyscan (Zeiss, Jena, Germany) by z-stacked tile-scanning.

### Stereomicroscopic image analysis (macro-scale)

All whole-skin images were analyzed with custom-build IJ1 macros in Fiji ImageJ^42^. Whole RAG mouse skins measured 40.4 cm^2^ (RAG15) to 82.5 cm^2^ (RAG10) in surface area (**Table S1**). In comparison, C57Bl.6 mouse skins are negligibly thin, measuring on average 0.034 cm (female) and 0.037 cm (male) in thickness^43^. Thus, we dismiss tissue thickness at the macro-scale and treated whole-skin images as two-dimensional surfaces to simplify the analysis. Patch detection was based on 8-bit grey-scale analysis of the *L. donovani* tdTomato-signal in the isolated red channel. The bright fluorescent signal of densely accumulated amastigotes per host cell strongly enhances the fluorescent-signal far beyond what endogenously fluorescent myeloid cells could have achieve. The obligatory intracellular nature of *Lesihmania* amastigotes allowed us to use the parasite signal as a proxy for infected myeloid cell, which allowed their indirect detection.

To determine a reliable detection threshold, we applied a three-level approach of the “Multi-Otsu-Threshold” algorithm plugin for Fiji ImageJ to tdTom-*L. donovani* infected and naïve RAG skins^44^. Naïve skins rendered maximum intensity thresholds <35, while infected skin showed a range of about 60-150 on an 8-bit grey-scale. For best detection sensitivity verses specificity balance, we chose a threshold of 70 on an 8-bit grey-scale. Further, we had to define the minimum size of a patch. Best image resolution allowed for a pixel size of 21.47 μm x 21.47 μm = 461 μm^2^, which did not allow to resolve for individual *L. donovani* amastigote, which are 2 – 3 μm in diameter (*r*^2^ * π = 1.5^2^ * 3.1416 ≈ 7.1 μm^2^). Within a mammalian host, *L. donovani* amastigotes are intracellular; most commonly residing in macrophages^45^. Thus, *L. donovani* amastigote distribution can be described by their host cell distribution. The cell body shape of macrophages and DCs is plastic and in tissues like the skin highly irregular, making it difficult to calculate the 2D surface area of such cells. Assuming that the overall cell surface is the same between a rounded and irregularly shaped macrophage, accepting an average diameter of 25 μm and assuming a flat cell body shape, we can calculate an approximate 2D cell area (*r*^2^ * π = 12.5^2^ * 3.1416 ≈ 491 μm^2^)^46^. A single macrophage is therefore defined by at least 4 pixels. We thereby defined the smallest arguable patch size as 2 x 2 macrophages (at least 9 pixels) in 2D. Thus, (3 * 21.47 μm)^2^ = 4148.65 μm^2^ was set as the smallest possible size to define patches in our stereomicroscopic images. Using these signal detection and patch size thresholds on naïve RAG skins showed false positive detection of ≤15 small patches (<0.1 mm^2^), which could largely be attributed to dust particles and other autofluorescent events (**Fig. S22A**). Further, manual patch counting in 1 cm^2^ areas on infected skins was used to assess the false negative detection rate, showing underestimation of very small size (<0.1 mm^2^; **Fig. S22B**) and loss of detection sensitivity at larger patch fringes, underestimating patch areas. The macro for density-peak detection was adapted from a publicly available protocol on particle counting in cells (Andrew McNaughton, University of Otago, NZ).

Further, two standard plugins available for Fiji ImageJ were used; a plugin published by Haeri & Haeri (2015) for nearest neighbor distance (NND) analysis^47^ and a plugin published by Ben Tupper to analyze patch clustering^48^. For the latter, a maximum distance of 3.9 mm between patch boarders was chosen. This value was based on the idea that an average sand fly is from proboscis tip to the abdominal posterior about 2 mm long. If we used just <2 mm (1.95 mm) as a radius to mark a circular landing zone for sand flies that landing zone has diameter of 3.9 mm (**Fig. S1D**). Conceptually, any patch reaching into the landing zone may be immediately accessible to a sand fly without moving away from its landing spot. Even if a parasite patches bordered just at the periphery of the landing zone, parasites from these patches will end up in the blood pool, which has an average diameter of ∼1 mm^11^.

### Confocal image analysis (micro-scale)

Volocity (V.6; Quorum Technologies, Laughton, UK), was used for amastigote to epidermis distance measurements. Individual amastigotes were detected by eroding the tdTomato-signal until signal segments coincided with separate sets of YOYO-1 labelled small nuclei and kinetoplasts. The epidermis was demarcated by consolidating all Brilliant Violet 421 signal of Cytokeratin-10 (CK-10) positive areas into a single mass under exclusion of skin appendages. Distances were measured by taking the shortest distance from the centroid of the tdTom amastigotes to the edge of the CK-10 positive band. To collate distance measurements from images from the same RAG mouse and to compare between RAG mice, the measurements required normalization due variations in skin thickness. We estimated the mean thickness of the epidermis, dermis and hypodermis, respectively, for each image by averaging at least three cross measurements in different places for each layer and then adjusted all measurements by the factor of difference between the mean skin layer thickness and the optical skin thickness as measured by Sabino et al. (2016)^43^. While a difference in sex did not significantly affect the macroscale analysis, at the micro-scale skin shows significant difference between sexes. While epidermal thickness is comparable between male and female C57Bl.6 mice, the dermis of a male C57Bl.6 mouse is ∼1.4 times thicker than that of a female. Conversely, the hypodermis of female C57Bl.6 mouse is ∼1.84 times thicker than that of a male. Thus male and female B6.*RAG2^-/-^* mice are not directly comparable.

StrataQuest (TissueGnostics, Vienna, Austria), was used to measure *L. donovani* amastigote co-localization in and distribution of Man^R+^ and CD11c^+^ cells in the skin. Amastigotes were detected by segmentation on the tdTomato signal and host nuclei were detected by segmentation on a mask of YOYO-1 signal minus tdTomato signal. Host nuclei were used as a seed and a mask expanded based on Brilliant Violet 421 signal of Man^R^ and Alexa Fluor 647 signal of CD11c to determine cell type. The epidermis mask was demarcated by consolidating Brilliant Violet 421 signal of Man^R^, Dylight 650 signal of CD11c and YOYO-1 signal areas below the epidermis were excluded by size. Tissue morphology was used to define masks for the dermis and hypodermis. The tissue and cell masks were used to determine the location of the amastigotes.

### Code availability

All Fiji ImageJ macros and R scripts used in this study were deposited in on GitHub (https://github.com/joedoehl/LeishSkinSPPM) and are freely available for download.

### Statistics

All statistics were conducted in GraphPad PRISM v.8, SPSS v.25^49^ or Rstudio v.1.3.959^50^ running R v.4.0.2^51^. Prior to test application, parametric test assumption (normality by Shapiro-Wilk Test; equal variance by Brown-Forsythe test; etc) were tested for all data sets. If these assumptions were not met even after data transformation, appropriate non-parametric tests were selected. In case of the Quadrat test, which is based on chi-squared, chi-squared was replaced by a Fisher’s exact test whenever squares of the grid applied to the window of observations had several squares with <5 or at least one square with 0 observations, which was the case for all RAG mice. Prior to statistical analysis, all percent datasets were arcsin transformed. Applied statistical tests are stated throughout the text and in corresponding figure legends. Harvested mouse skins varied considerably in size between RAG mice (40.4 cm^2^ – 82.5 cm^2^). For comparison and correlation analyses between RAG mice, some data, like total patch area, patch counts, parasite counts were normalized to area / cm^2^ and counts / cm^2^, respectively. Where data skewness was an issue, we used the median as a more robust location measure. Spatial point pattern analysis was performed in Rstudio based on the work of Baddeley and co-workers^16^. The following R packages were used for code execution: abind^52^, deldir^53^, e1701^54^, ggpubr^55^, goftest^56^, graphics^51^, grDevices^51^, polyclip^57^, RandomFields^58^, rlist^59^, rms.gof^60^, scales^61^, sp^62^, sparr^63^, spatstat^16^, spatstat.utils^64^, stats^51^, tensor^65^, tidyverse^66^ and utils^51^. Each observation in the analysis was verified by at least two different tests. Running multiple test increases confidence in the result as different tests are not always in agreement.

### Spatial Point Patterns Methodology

A “spatial point pattern” is a dataset containing the spatial locations of events or objects as points within a window of observation. The characterization of the spatial arrangement of these points is the main focus of spatial point pattern methodologies. This is generally done by measuring inter-point distances and characterizing the dispersion between points. Such methodologies find application in a myriad of fields, including epidemiology, revealing important spatial features not discernable by eye^67^. Of course, not all “point data” permits analysis as a “point pattern”, particularly when sampling location is not relevant to the ultimately studied sample property. Conversely, datasets may represent point patterns which are not immediately recognizable, e.g. when the objects’ physical size exceeds technical classification as points; such limitations can be accommodated by choosing the coordinates of the center of mass of the objects as the point and attaching the objects’ size as an attribute, termed mark, to the respective points^68^. This was used in our study where the size of some observed parasite skin patches clearly exceeds the definition of point objects. Very large skin parasite patches also showed great heterogeneity in tdTomato signal density, containing multiple high-density areas. This suggested that very large skin patches were conglomerates of smaller patches that had coalesced forming continuous, although heterogeneously densely parasitized, patches.

We used functions provided by the spatstat R package by Baddeley and co-workers^16^. The following definitions are all taken from their work “*Spatial Point Patterns: Methodology and Applications with R*”^16^. Several functions used in our study were derived from the empirical K-function, originally proposed by Ripley in 1977^69^:

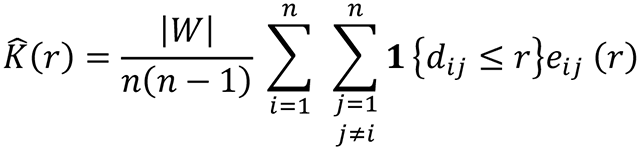

where *d_ij_* is the distance between all ordered paired points *x_i_* and *x_j_* (*d*_*ij*_ = ❘|*x*_*i*_ − *x*_*j*_|❘), *r* the value of measured distances (*r* ≥ 0), |*W*| the area of the observation window, *n*(*nn* − 1) the total number pairs of distinct points, 1{…} the ‘indicator’ notation, which is 1 when statement ‘…’ is true and 0 when false and *e*_*ij*_(*r*) an edge correction weight. Riley’s K-function implicitly assumes that the investigated point process (***X***) has homogeneous intensity and is stationary. If a point process is suspected to be inhomogeneous, then that must be considered in the calculation of the K-function. The standard estimator of *K* can be extended to produce the inhomogeneous K-function:

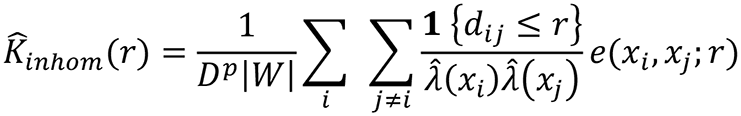

where *e*(*x*_*i*_, *x*_*j*_; *r*) is an edge correct weight and 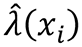 *and* 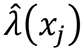 are estimates of the intensity functions *λ*(*x*_*i*_) *and λ*(*x*_*j*_), respectively. The constant *D*^*p*^ is the *p*^th^ power of

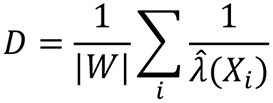

The inhomogeneous K-function assumes that the spatial scale of interaction remains constant, while the intensity is spatially varying. However, the spatial scale may be locally dependent. In this case, a rescaled template process may be applied that is locally stationary and isotropic, but shows a varying rescaling factor between locations within the observation window. Thus, the empirical K-function is adjusted for each pair of data points (*x*_*i*_, *x*_*j*_) by calculating the rescaled distance

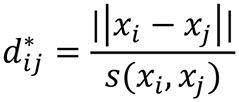

where the rescaling factor is

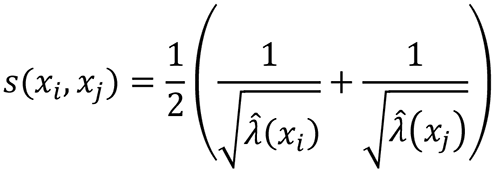

This means the locally scaled K-function is defined by

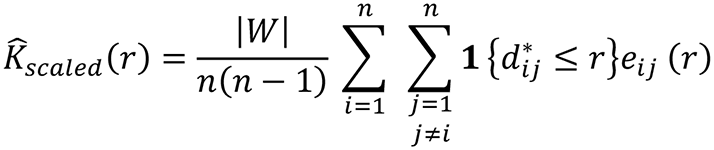

The L-function, as proposed by Besag^69^, is then derived from the various K-function by

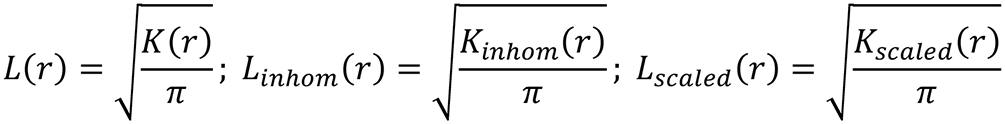

respectively. The pair correlation function is derived from the K-function as

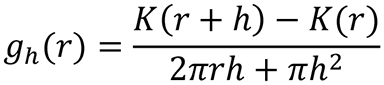

When *h* becomes very small, then *π*ℎ^2^ becomes negligible and 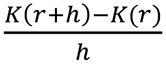becomes the derivative of the K-function with respect to *r* (*K’*(*r*)). Thus, in two dimensions, the pair correlation function can be defined by

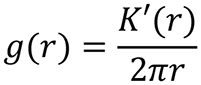

More details can be found in Baddeley & co-workers book “*Spatial Point Patterns: Methodology and Applications with R*”^16^.

### Data availability

The authors declare that the data supporting the findings of the study are available within the article and its Supplementary Information files or are available from the authors upon request.

## Acknowledgments

This work was supported by a Wellcome Senior Investigator Award (# WT106203; http://www.wellcome.co.uk) and a MRC Programme Grant (#G1000230; http://www.mrc.ac.uk) to P.M.K. The authors thank Andrew McNaughton from the Confocal / µCT Facility at University of Otago, NZ, for the permission of adapting his protocol and invaluable discussion, Jesus Valenzuela from the VMBU-LMVR at the NIAID/NIH, USA, for encouragement and time made available to complete the manuscript, the Imaging and Cytometry lab and staff, in particular Graeme Park and Jo Marrison, in the Bioscience Technology Facility at the University of York, UK, for use of their equipment and technically knowhow that supported the image acquisition and the staff of the Biological Services Facility at the University of York, UK, for animal husbandry.

## Author contributions

J.S.P.D. and P.M.K. designed the study; J.S.P.D., H.A., N.B. and A.R. conducted the experimental studies; J.S.P.D. conducted the spatial point pattern analysis, modelling and all the statistical analysis, which were revised by S.C. and J.W.P.; the manuscript was written by J.S.P.D. and P.M.K. and all authors contributed to the final version; P.M.K. supervised the study and has senior authorship; P.M.K. conceived the study.

## Competing financial interest

There are no competing financial interests in relation to this study.

## Notes

### Competing Interest Statement

The authors have declared no competing interest.

### Summary of Updates

Minor text revisions; enhanced description of statistical and modelling methodologies.

