## Supplemental Figures 1-22 and Tables 1-3 for "Spatial point pattern analysis identifies mechanisms shaping the skin parasite landscape in *Leishmania donovani* infection"

**This file includes:**

Supplementary Figure Legends

Supplementary Figures 1 to 22

Supplementary Table 1 to 3

### Supplementary Figure Legends

**Figure S1 | Schematics of experimental designs and methodologies**

(A) Experimental schematic representing the treatment of RAG1-10 & RAG15-18. (B) Experimental schematic representing the treatment of RAG11-14 (EXP5) & RAG19-23 (EXP7). (C) Schematic representation of technique for mouse skinning. (D) Schematic representation of the concept for the nearest-neighbor distance pattern analysis.

**Figure S2 | Graphs from figure 1 resolved by individual RAG mouse**

(A) Quantitative PCR data of estimated parasite loads of 24 or 16 biopsies (ø 0.6 cm) / RAG mouse. This graph shows the same data as in Fig. 1A resolved by individual RAG mice (Kruskal-Wallis Test: P<0.0001). (B) Skin parasite patch area data from Fig. 1B resolved by respective RAG mice (Kruskal-Wallis Test: P<0.0001). (C) Mean parasite patch density / patch as shown in Fig. 1F resolved by respective RAG mice (Kruskal-Wallis Test: P<0.0001).

**Figure S3 | 23 RAG mouse skin composite images showing detected parasite patches**

Composite stereomicroscopic image of the *L. donovani* tdTomato-signal as described in the methods section. All detected skin parasite patches in the ImageJ analysis are framed by yellow outlines. Note: The Photoshop composite image for RAG2 and RAG14 caused an irreversible background error in the green channel when read into ImageJ, giving these two images a green hue in the output. However, all data acquisition was done in the red channel and thus, this error had no effect on the data collection

**Figure S4 | Bar graphs from figure 1 E resolved by individual RAG mouse**

(A) Bar chart of the ratio of patch counts / size category showing the same data as in Fig.1E ‘A’ resolved by the respective RAG mice (Friedman Test: P=0.7544). (B) Bar chart of the contribution of size category to the overall patch coverage showing the same data as in Fig.1E ‘B’ resolved by the respective RAG mice (Friedman Test: P=0.7345).

**Figure S5 | Schematic distribution of skin parasite patches marked by patch area**

Skin parasite patches were categorized by patch size according to the log-scale employed in Fig.1E and plotted according to the location of the center of mass for each skin parasite patch.

**Figure S6 | Pixel density skin images**

8-bit grey-scale images, like in Fig.1H, showing the tdTomato light intensity per pixel for each analyzed RAG mouse, respectively. Dark blue refers to lowest light intensity, while white refers to highest light intensity. See scale for reference. In magenta are the outlines of detected skin parasite patches.

**Figure S7 | Representation of skin parasite patches as spatial point pattern without marks**

Each red circle marks the center of mass of parasite patches within the skin. The circle size does not convey any measure here.

**Figure S8 | Representation of skin parasite patches as spatial point pattern with area mark**

Each red circle marks the center of mass of parasite patches within the skin. The circle size corresponds to the relative area of each patch without information on the shape of the patch. The circle size scale is specific to each graphical representation.

**Figure S9 | Intensity measurement plots without marks**

The images show how closely skin parasite patches are clustered together in different sights of the skin. These intensity measurements do not take account of patch size. On the color scale, blue-purple colors indicate low skin parasite patch density, while orange-yellow indicate high skin parasite patch density. The scale is relative to each image.

**Figure S10 | Intensity measurement plots with area mark**

The images show the distribution of skin parasite patches by patch area. On the color scale, blue-purple colors indicate the presence of skin parasite patch with small patch area, while orange-yellow indicate the presence of skin parasite patch with large patch area. The scale is relative to each image.

**Figure S11 | Graphical representation of cluster analysis**

The red lines indicate where patches are separated ≤3.9 mm from patch periphery to patch periphery. Individual patches are marked in different shadings of grey. The red lines indicate where networks of closely placed skin parasite patches may enhance the likelihood of biting sand fly finding a patch, while the shaded areas represent the patches.

**Figure S12 | Test for isotropy of skin parasite patch point patterns**

The red dotted lines mark the theoretical homogeneous Poisson process as a null hypothesis. The black solid lines represent the plotted data by distance of patches in mm (*r*). The grey shaded area around the solid black line is the 95% confidence interval. Should the red dotted line be not included into the grey shaded area at any stage then the point pattern is not isotropic. RAG14 did not produce a result due to too few skin parasite patches for the analysis.

**Figure S13 | Correlation-stationary test using the inhomogeneous K-function**

The double graphs show the theoretical inhomogeneous Poisson process as a null hypothesis and up to four graphs based on different border correction approaches. Please, consult the respective graph legends for the color code of each graph.

**Figure S14 | Correlation-stationary test using the locally scaled K-function**

The double graphs show the theoretical inhomogeneous Poisson process as a null hypothesis and up to four graphs based on different border correction approaches. Please, consult the respective graph legends for the color code of each graph.

**Figure S15 | Scaled L-function of skin parasite patch distribution**

The red dotted line represents the theoretical locally scaled Poisson process as a null hypothesis. The solid black line shows the plotted data. The data is presented against the distance '*r*' between skin parasite patches measured in mm. The grey areas are simulation-envelopes around the theoretical Poisson process. Departure of the solid black line above or below the red dotted line are indicative of clustered and regular distributions instead of an independent distribution, respectively. Departure of the solid black line from the grey shaded simulation-envelops indicates significant departure from the theoretical Poisson process.

**Figure S16 | Pair correlation function plots**

The red dotted line represents the theoretical locally scaled Poisson process as a null hypothesis. The solid black line shows the plotted data. Departure of the solid black line above or below the red dotted line are indicative of clustered and regular distributions instead of an independent distribution, respectively.

**Figure S17 | K-function of point process residuals**

The red dotted line represents the theoretical locally scaled Poisson process as a null hypothesis. The solid black line shows the plotted data. The blue and green line mark the limits of the upper and lower confidence interval. Departure of the solid black line above or below the red dotted line indicates significant departure of data point independence. Further, departure of the solid black line above or below the red dotted line are indicative of clustered and regular distributions instead of an independent distribution, respectively.

**Figure S18 | G-function of point process residuals**

The red dotted line represents the theoretical locally scaled Poisson process as a null hypothesis. The solid black line shows the plotted data. The blue and green line mark the limits of the upper and lower confidence interval. Departure of the solid black line above or below the red dotted line indicates significant departure of data point independence. Further, departure of the solid black line above or below the red dotted line are indicative of clustered and regular distributions instead of an independent distribution, respectively.

**Figure S19 | Scaled L-function of simulated skin parasite patch distribution**

The red dotted line represents the theoretical locally scaled Poisson process as a null hypothesis. The solid black line shows the plotted data. The data is presented against the distance ‘r’ between skin parasite patches measured in mm. The grey areas are simulation-envelopes around the theoretical Poisson process. Departure of the solid black line above or below the red dotted line are indicative of clustered and regular distributions instead of an independent distribution, respectively. Departure of the solid black line from the grey shaded simulation-envelops indicates significant departure from the theoretical Poisson process.

**Figure S20 | Simulation output according to best fit model**

Graphical output of four iterations of the respective best-fit model simulation per RAG mouse.

**Figure S21 | Visual summary GIF of the proposed mechanism for skin parasite patch dispersal**

After an initial *Leishmania* amastigote seeding event in the host skin by arrival of infected host cells presumably from circulation, uninfected phagocytic cells in skin are attracted to the infected cells to form a type of innate granuloma. Attracted naïve phagocytic cells are then gradually infected in the innate granuloma, forming a parasite patch. As parasite patches keep growing, they seed new patches within a radius ‘r’ around themselves, potentially by escape of infected host cells or amastigote transfer through open-ended tunneling nanotubes. This process then keeps repeating itself, forming self-propagating networks of parasite patch clusters. These tightly spaced cluster networks may enhance the probability of naïve biting sand flies finding a dense parasite patch. All images were created with BioRender.com.

**Figure S22 | False positive and negative skin parasite patch detection analysis**

A) Naïve RAG mouse skin images were acquired and analyzed in the same fashion as positive skins to account for false positive detection. At close examination, detected event were identified as very small autofluorescent dust particles and other. Green squares denote the areas of enlargement in the naïve skin. B) Manuel skin parasite patch detection in 1 $cm^2$ areas in positive skin (marked by green squares) showed significant underestimation of small parasite patches (<0.1 $mm^2$). Yellow lines denote parasite detected by the ImageJ macro and green dotted circles indicated areas that were manually selected as potentially being true tdTomato fluorescence and, therefore, undetected skin parasite patches.

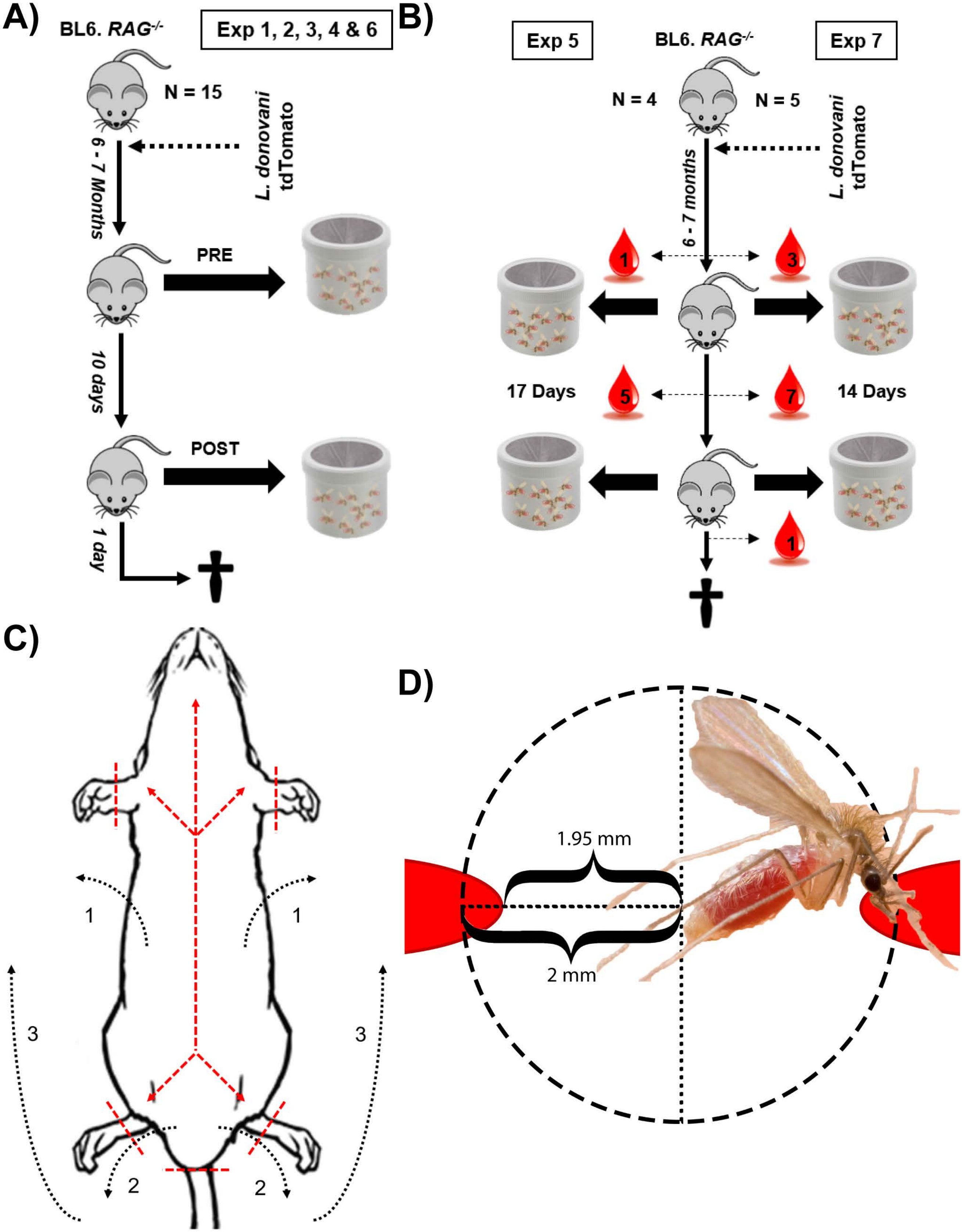
A)
BL6. RAG-/-
Exp 1, 2, 3, 4 & 6
N = 15
L. donovani tdTomato
6 - 7 Months
PRE
10 days
POST
1 day
B)
Exp 5
BL6. RAG-/-
Exp 7
N = 4
N = 5
L. donovani tdTomato
6 - 7 months
1
3
17 Days
5
7
14 Days
1
C)
1
1
3
3
2
2
D)
1.95 mm
2 mm

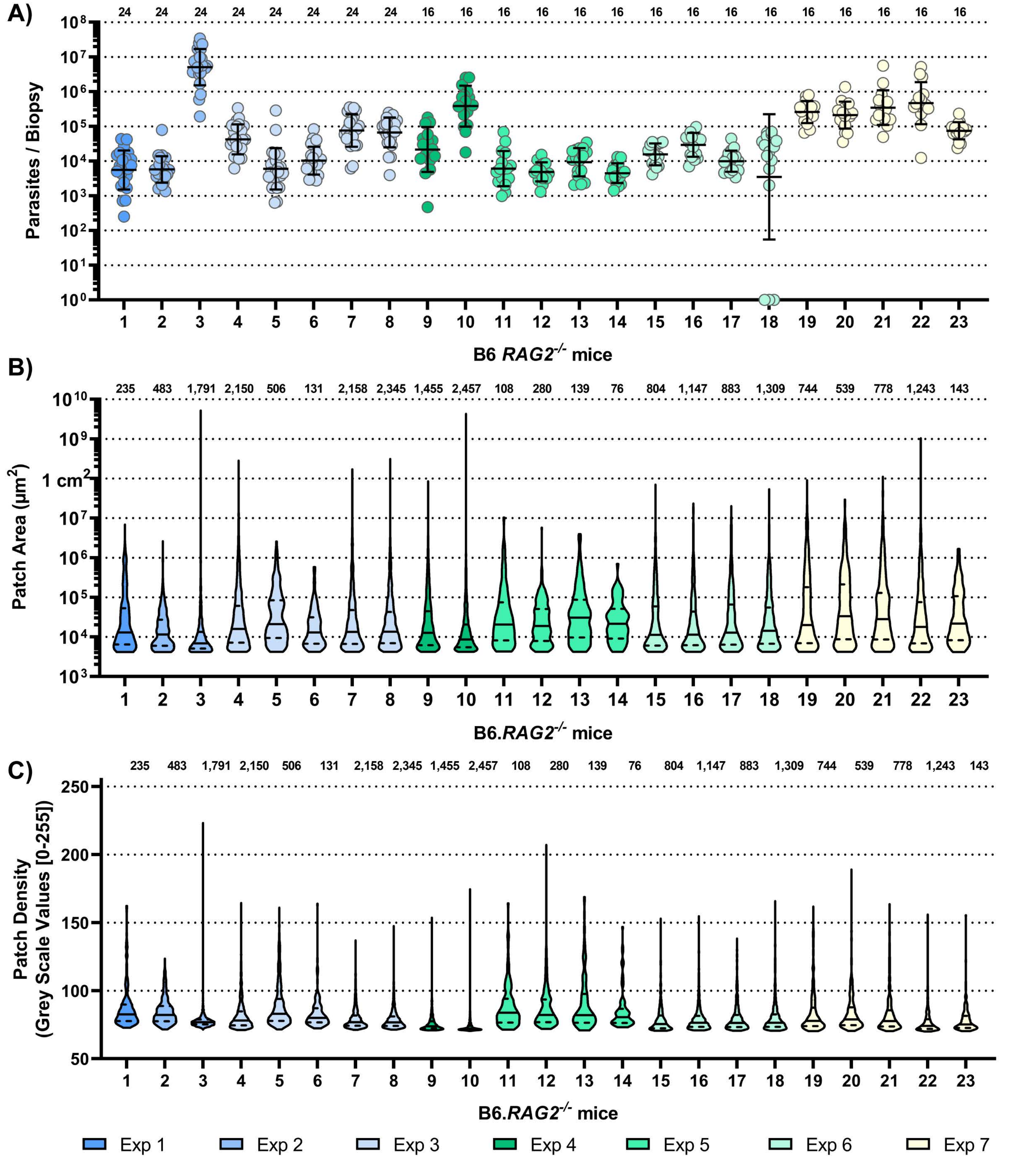
A)
B)
C)
Parasites / Biopsy
B6 RAG2-/- mice
Patch Area (µm2)
1 cm2
B6.RAG2-/- mice
Patch Density
(Grey Scale Values [0-255])
B6.RAG2-/- mice
Exp 1
Exp 2
Exp 3
Exp 4
Exp 5
Exp 6
Exp 7

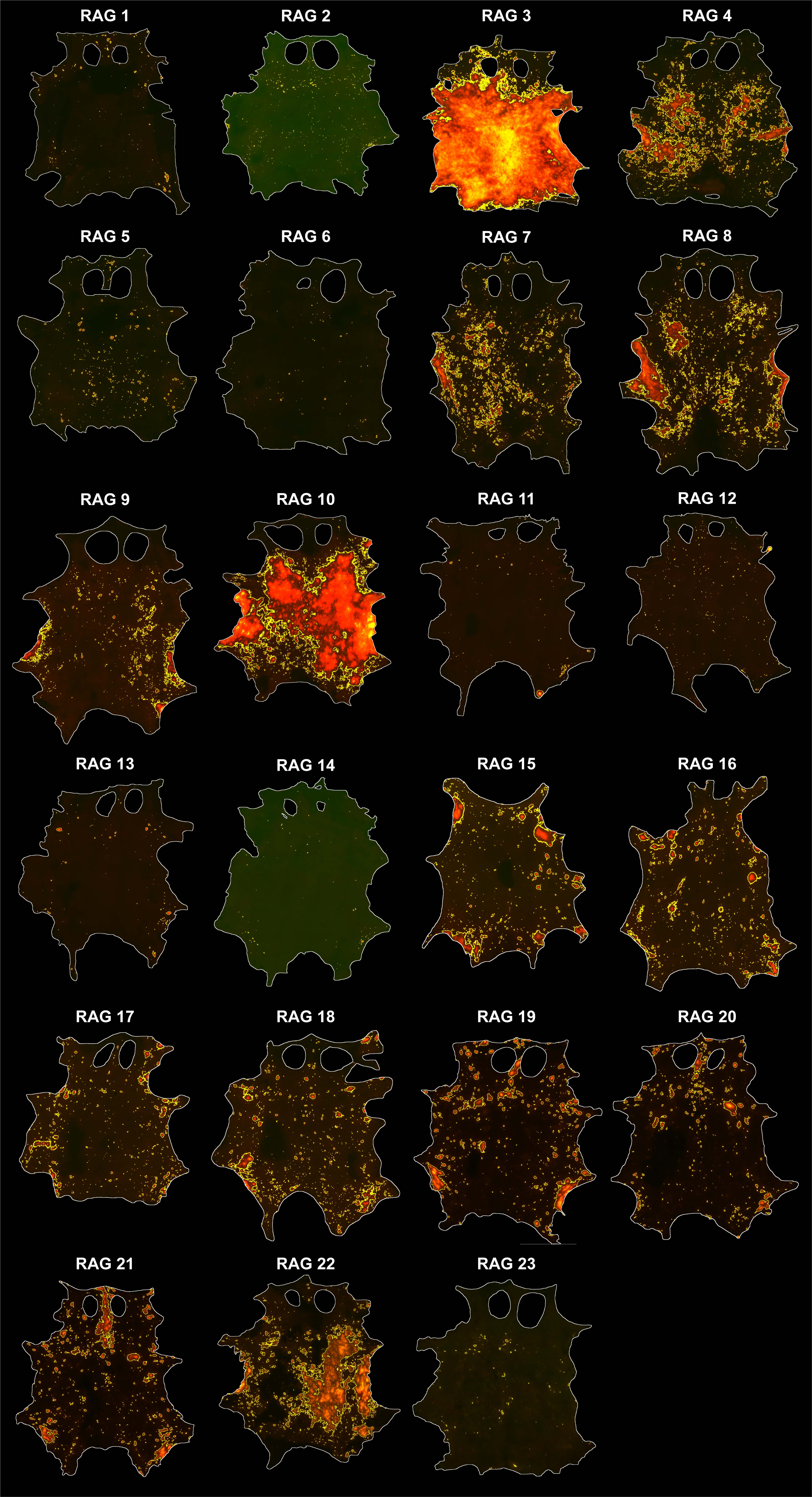

RAG 1
RAG 2
RAG 3
RAG 4
RAG 5
RAG 6
RAG 7
RAG 8
RAG 9
RAG 10
RAG 11
RAG 12
RAG 13
RAG 14
RAG 15
RAG 16
RAG 17
RAG 18
RAG 19
RAG 20
RAG 21
RAG 22
RAG 23

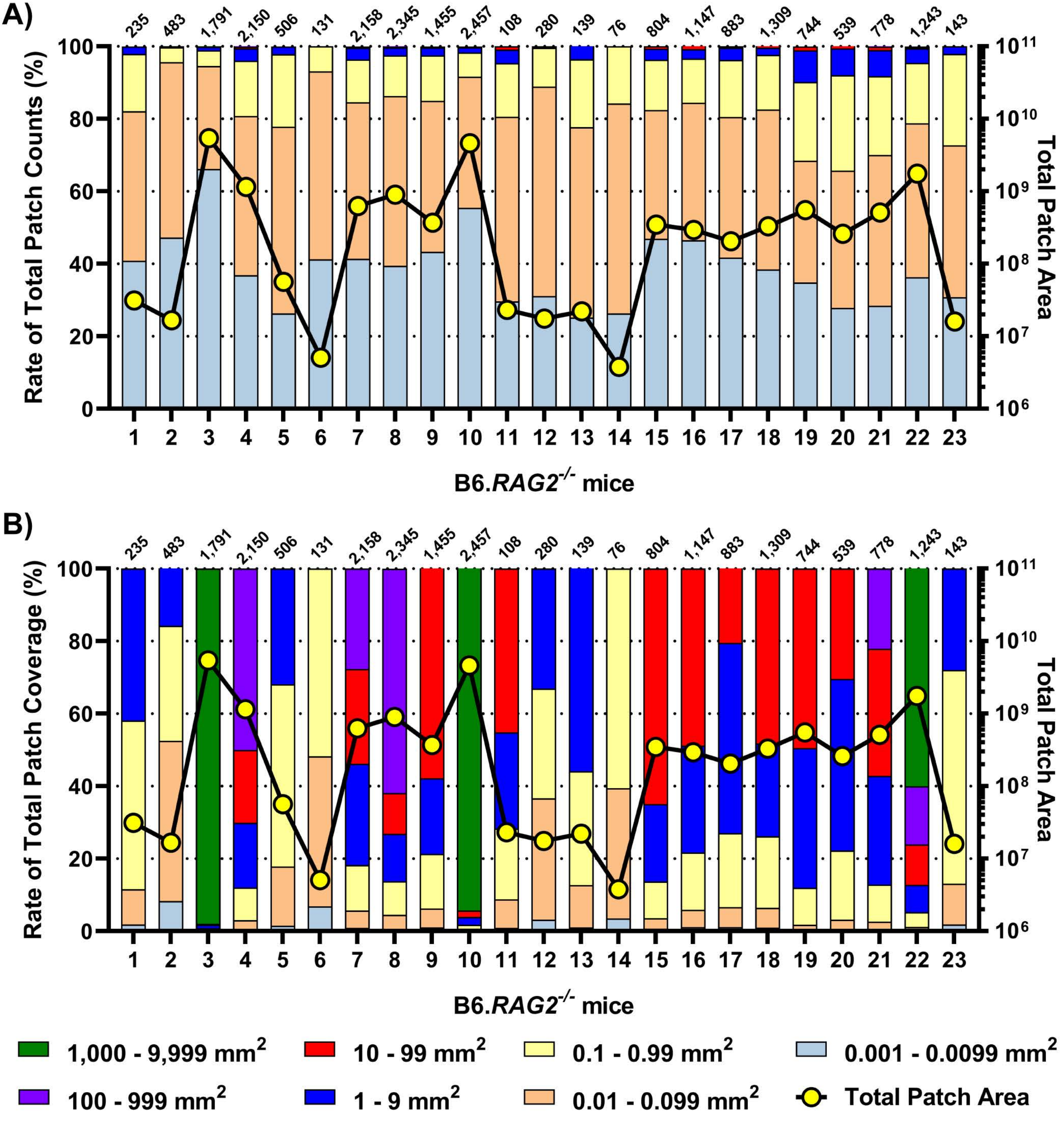
A)
235 483 1,791 2,150 506 131 2,158 2,345 1,455 2,457 108 280 139 76 804 1,147 883 1,309 744 539 778 1,243 143
Rate of Total Patch Counts (%)
Total Patch Area
B6.RAG2-/- mice
B)
235 483 1,791 2,150 506 131 2,158 2,345 1,455 2,457 108 280 139 76 804 1,147 883 1,309 744 539 778 1,243 143
Rate of Total Patch Coverage (%)
Total Patch Area
B6.RAG2-/- mice
1,000 - 9,999 mm2
10 - 99 mm2
0.1 - 0.99 mm2
0.001 - 0.0099 mm2
100 - 999 mm2
1 - 9 mm2
0.01 - 0.099 mm2
Total Patch Area

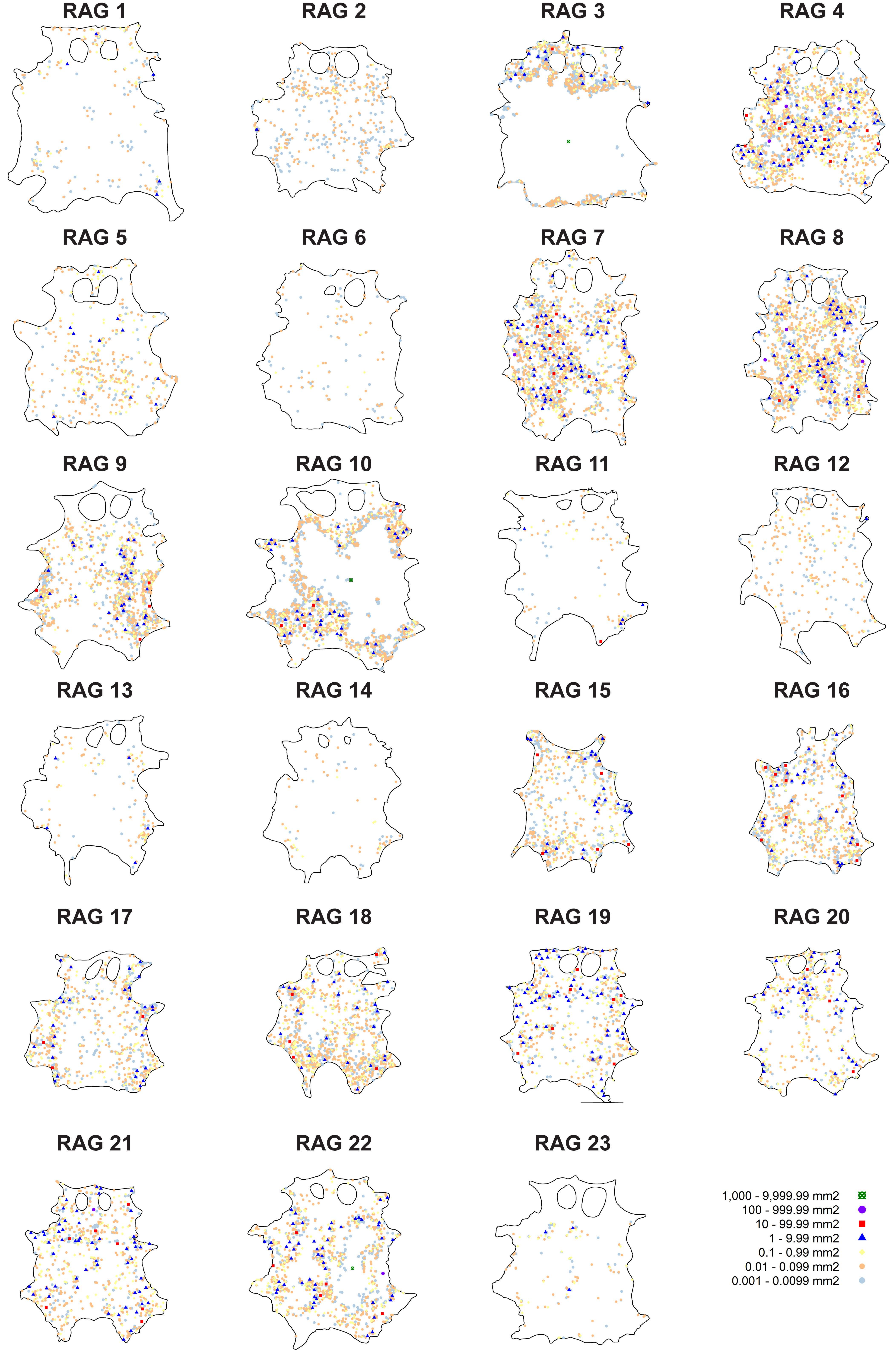
RAG 1
RAG 2
RAG 3
RAG 4
RAG 5
RAG 6
RAG 7
RAG 8
RAG 9
RAG 10
RAG 11
RAG 12
RAG 13
RAG 14
RAG 15
RAG 16
RAG 17
RAG 18
RAG 19
RAG 20
RAG 21
RAG 22
RAG 23
1,000 - 9,999.99 mm2
100 - 999.99 mm2
10 - 99.99 mm2
1 - 9.99 mm2
0.1 - 0.99 mm2
0.01 - 0.099 mm2
0.001 - 0.0099 mm2

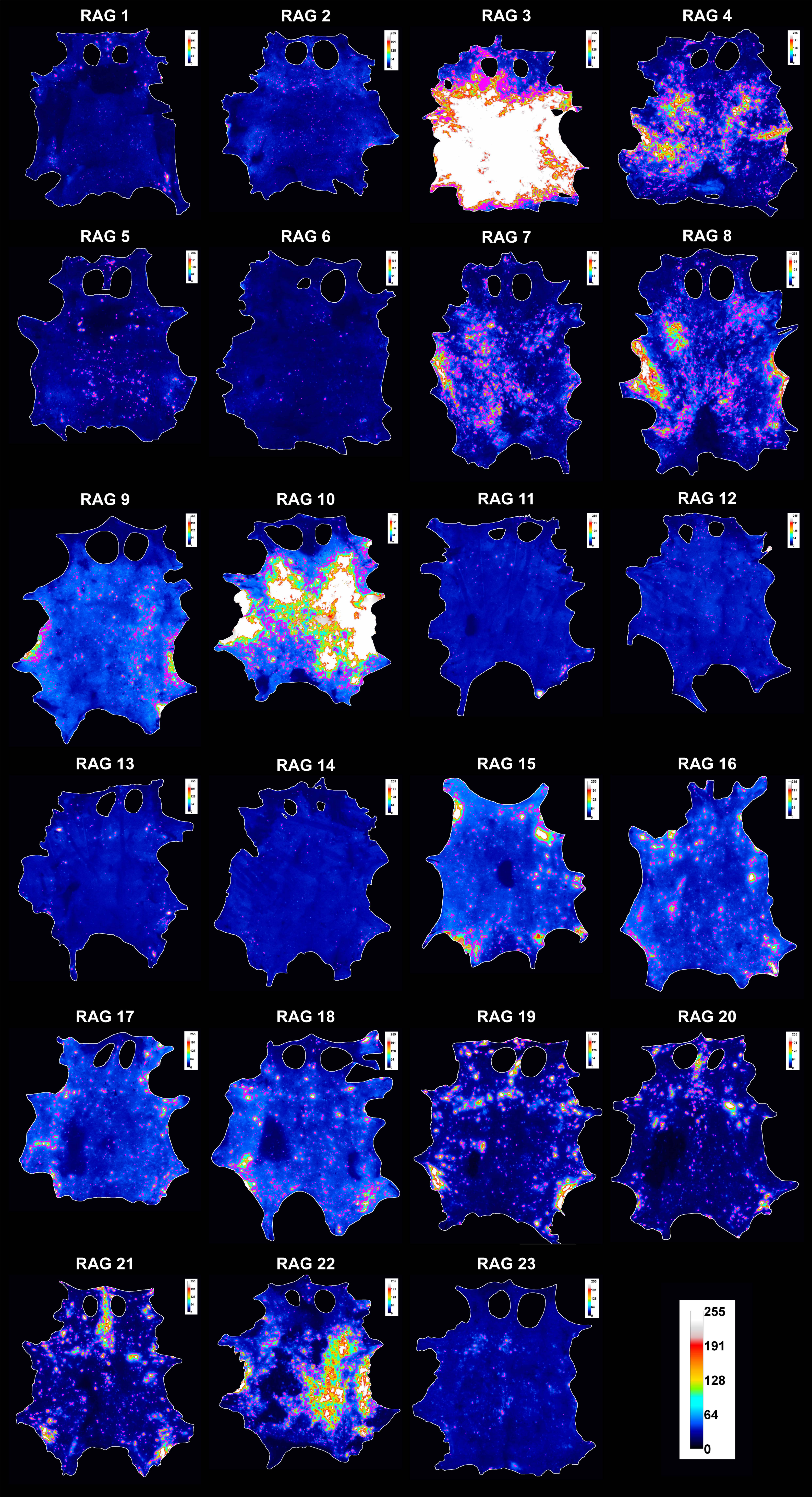
RAG 1
RAG 2
RAG 3
RAG 4
RAG 5
RAG 6
RAG 7
RAG 8
RAG 9
RAG 10
RAG 11
RAG 12
RAG 13
RAG 14
RAG 15
RAG 16
RAG 17
RAG 18
RAG 19
RAG 20
RAG 21
RAG 22
RAG 23
255
191
128
64
0

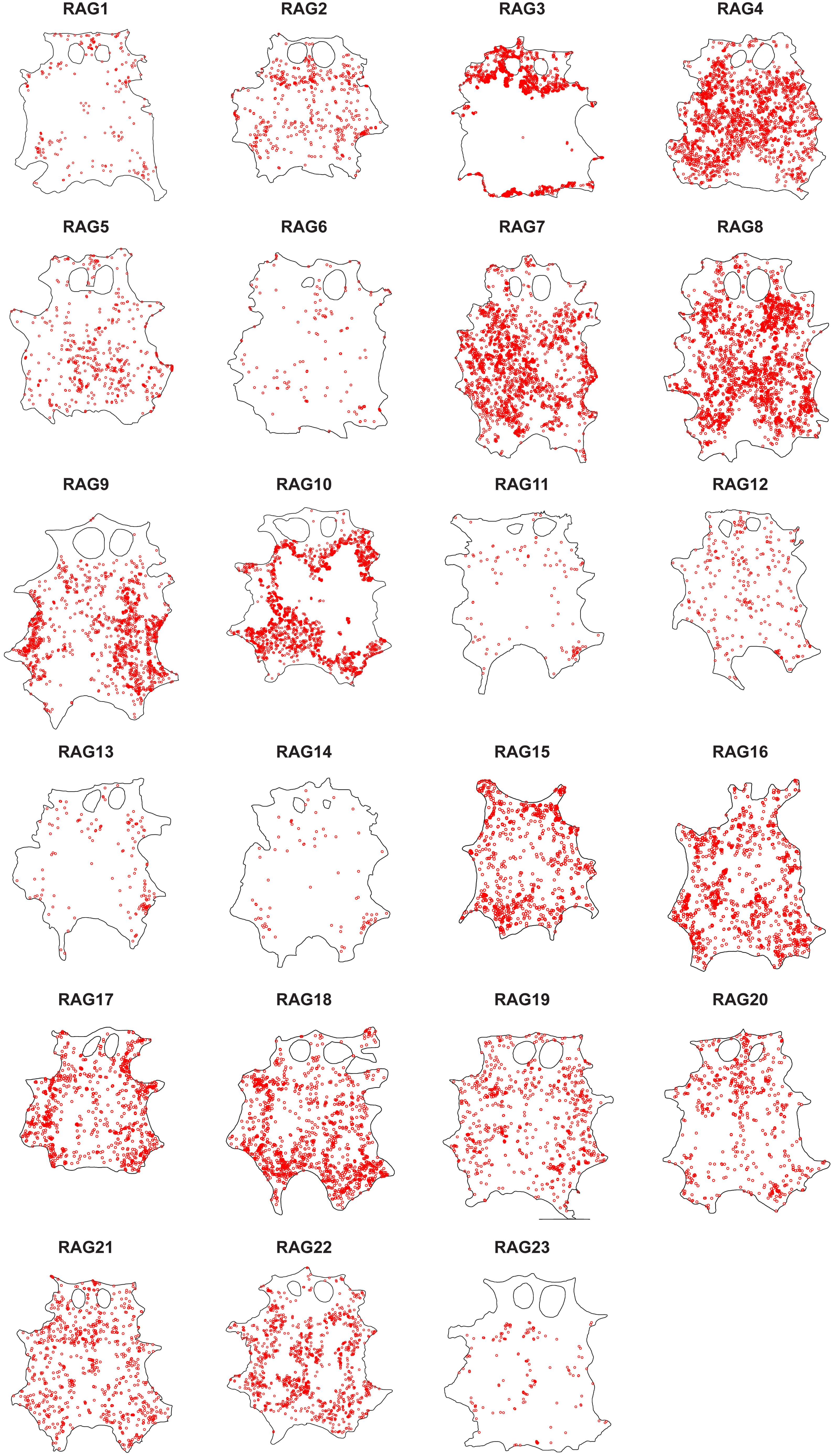
RAG1
RAG2
RAG3
RAG4
RAG5
RAG6
RAG7
RAG8
RAG9
RAG10
RAG11
RAG12
RAG13
RAG14
RAG15
RAG16
RAG17
RAG18
RAG19
RAG20
RAG21
RAG22
RAG23

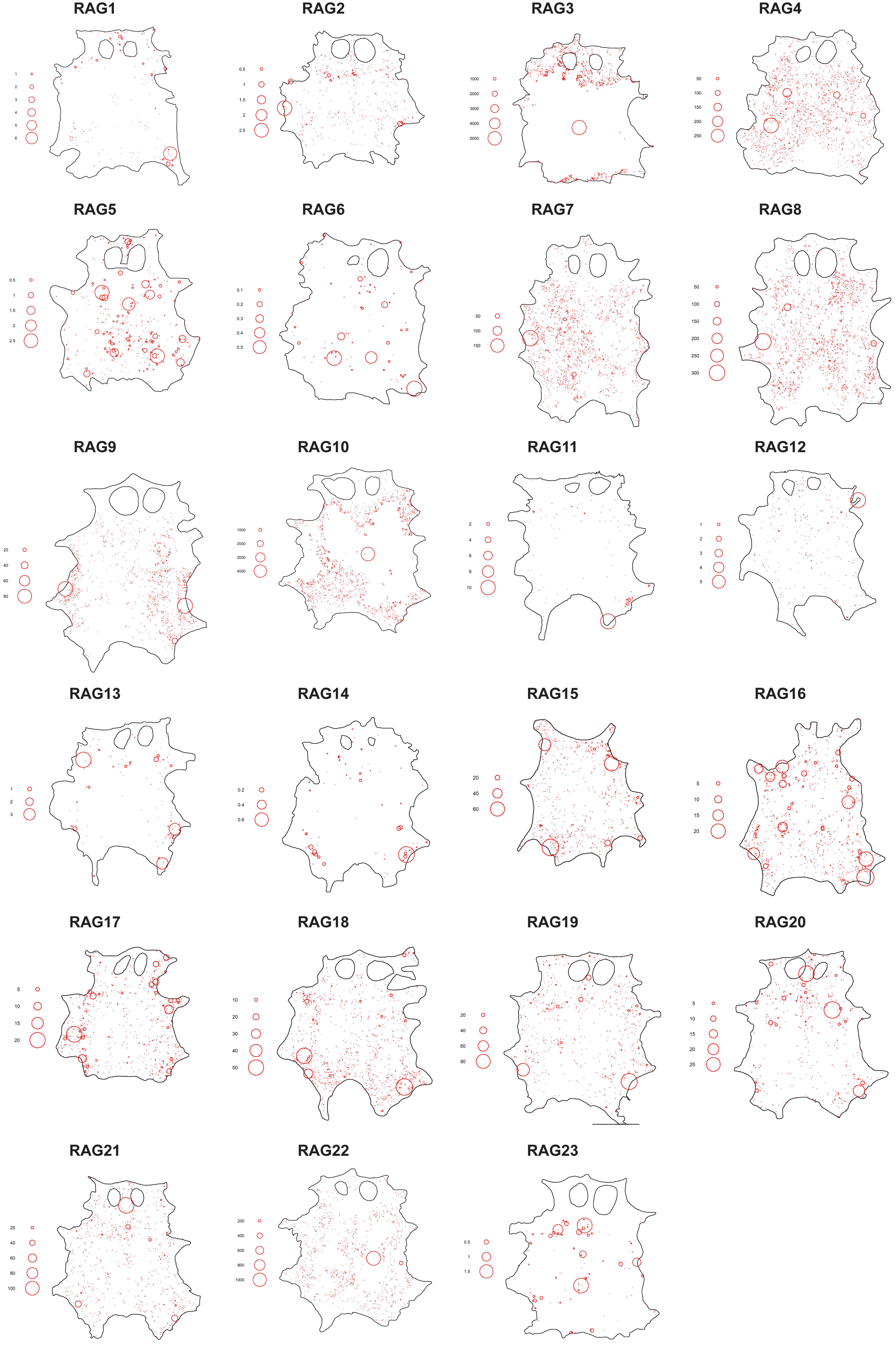
RAG1
1
2
3
4
5
6
RAG2
0.5
1
1.5
2
2.5
RAG3
1000
2000
3000
4000
5000
RAG4
50
100
150
200
250
RAG5
0.5
1
1.5
2
2.5
RAG6
0.1
0.2
0.3
0.4
0.5
RAG7
50
100
150
RAG8
50
100
150
200
250
300
RAG9
20
40
60
80
RAG10
1000
2000
3000
4000
RAG11
2
4
6
8
10
RAG12
1
2
3
4
5
RAG13
1
2
3
RAG14
0.2
0.4
0.6
RAG15
20
40
60
RAG16
5
10
15
20
RAG17
5
10
15
20
RAG18
10
20
30
40
50
RAG19
20
40
60
80
RAG20
5
10
15
20
25
RAG21
20
40
60
80
100
RAG22
200
400
600
800
1000
RAG23
0.5
1
1.5

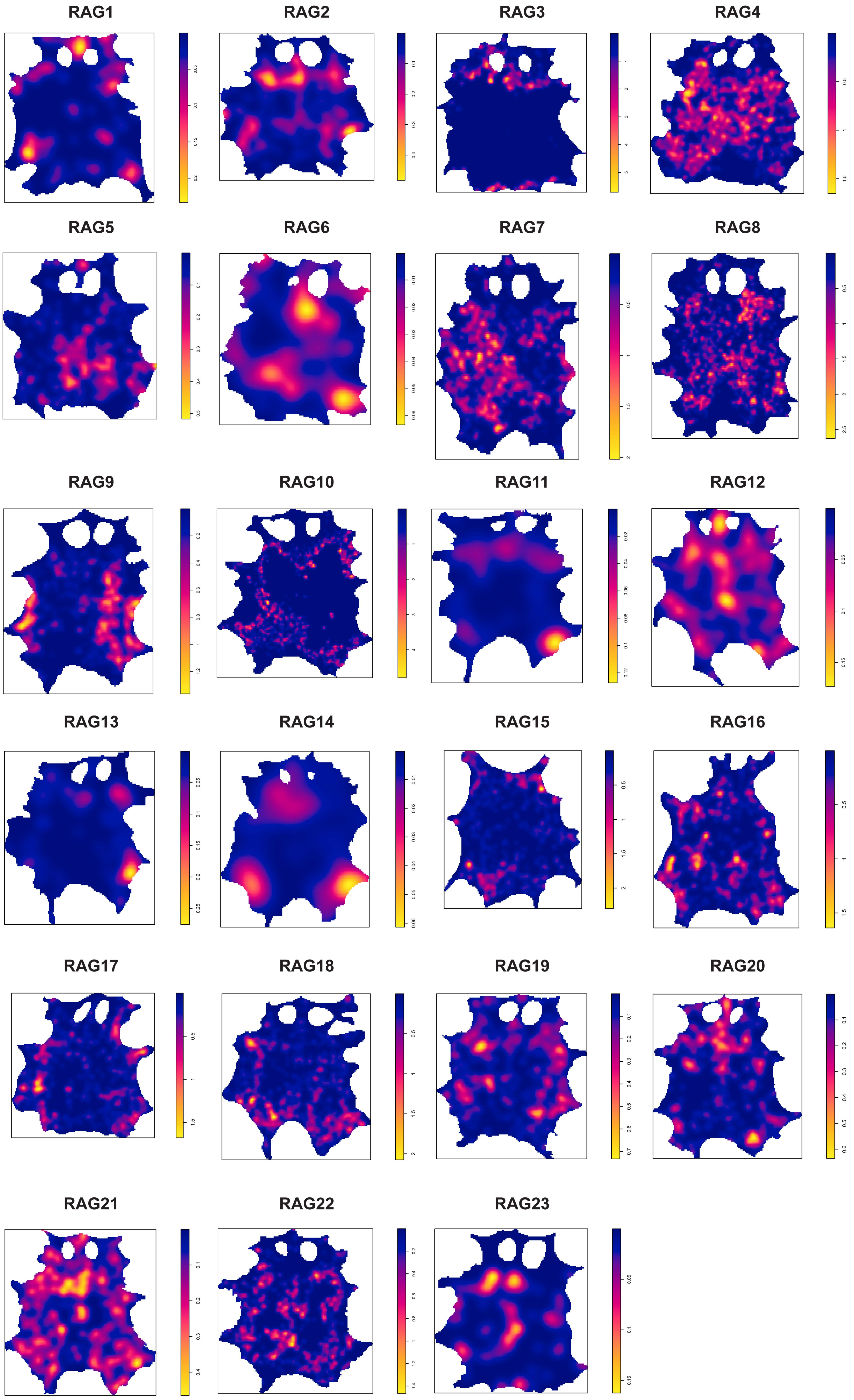
RAG1
RAG2
RAG3
RAG4
RAG5
RAG6
RAG7
RAG8
RAG9
RAG10
RAG11
RAG12
RAG13
RAG14
RAG15
RAG16
RAG17
RAG18
RAG19
RAG20
RAG21
RAG22
RAG23

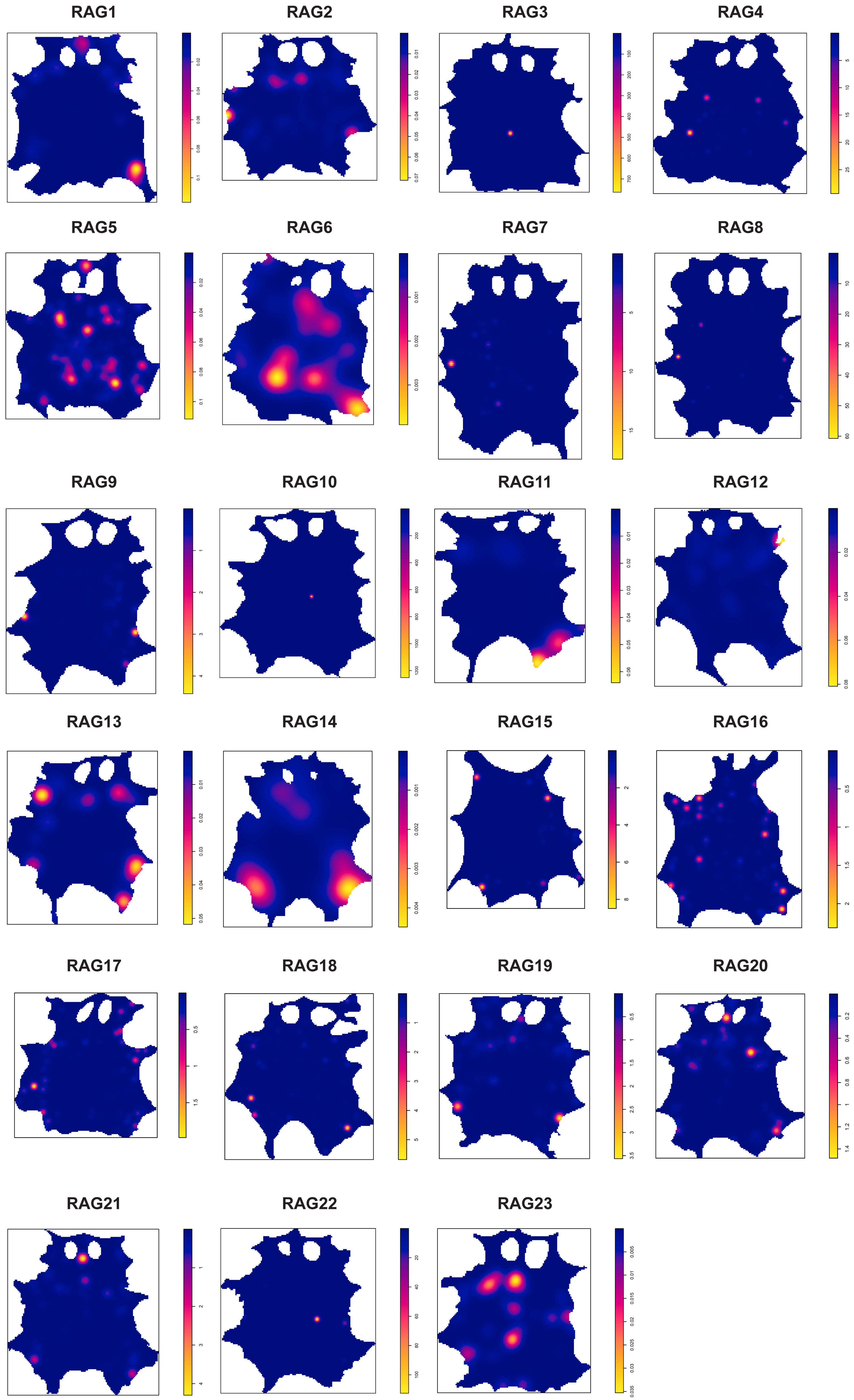
RAG1
RAG2
RAG3
RAG4
RAG5
RAG6
RAG7
RAG8
RAG9
RAG10
RAG11
RAG12
RAG13
RAG14
RAG15
RAG16
RAG17
RAG18
RAG19
RAG20
RAG21
RAG22
RAG23

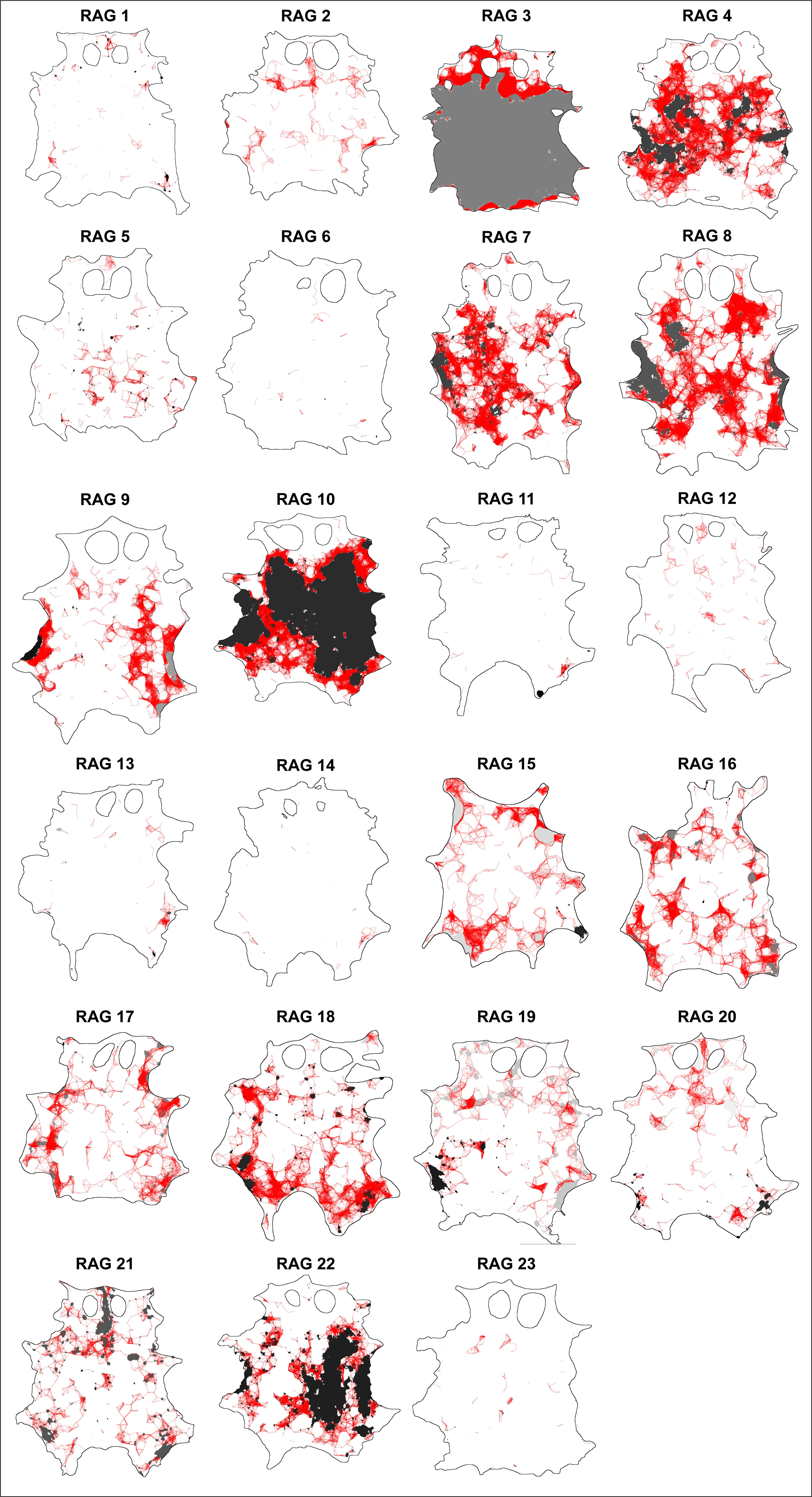
RAG 1
RAG 2
RAG 3
RAG 4
RAG 5
RAG 6
RAG 7
RAG 8
RAG 9
RAG 10
RAG 11
RAG 12
RAG 13
RAG 14
RAG 15
RAG 16
RAG 17
RAG 18
RAG 19
RAG 20
RAG 21
RAG 22
RAG 23

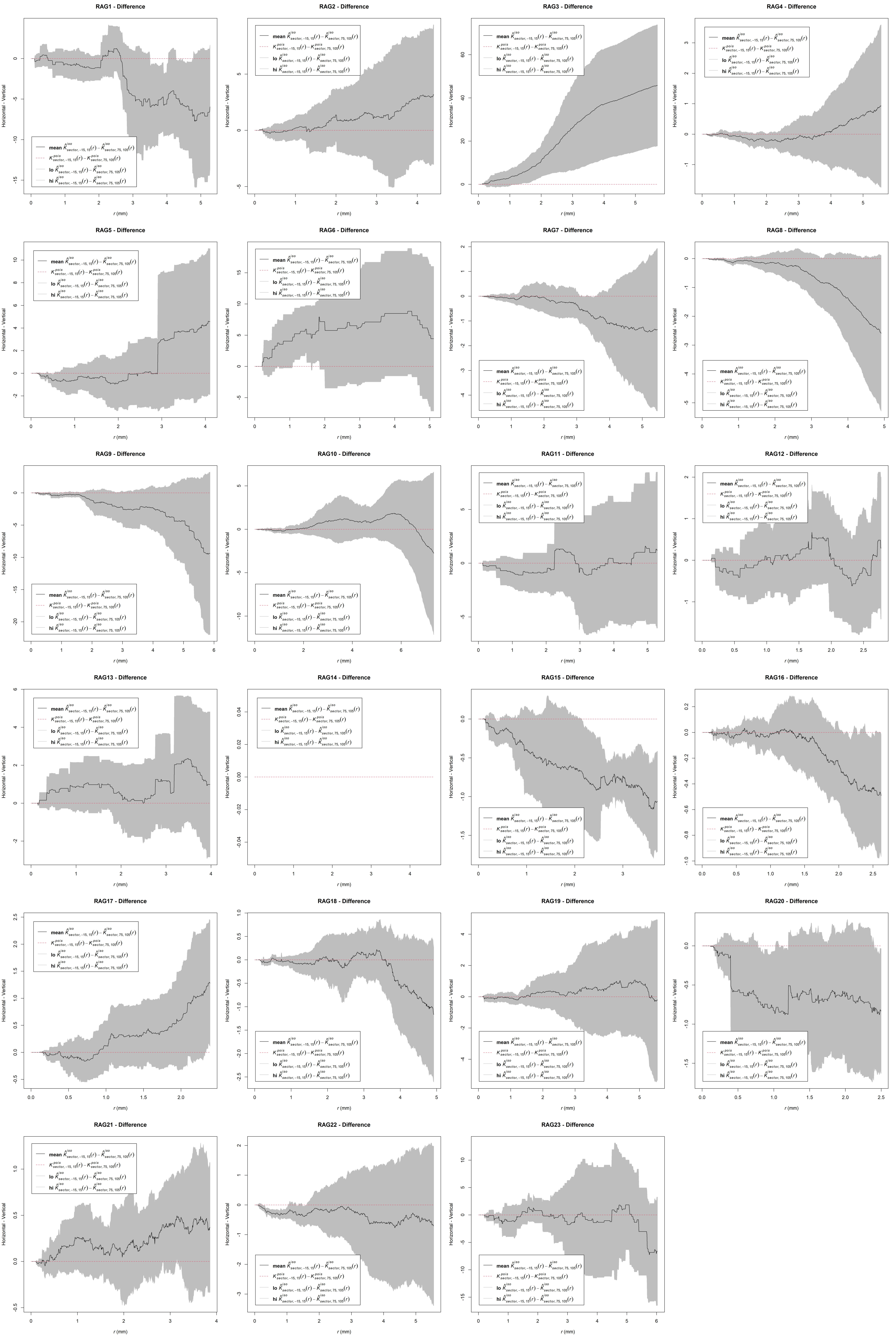
RAG1 - Difference
RAG2 - Difference
RAG3 - Difference
RAG4 - Difference
RAG5 - Difference
RAG6 - Difference
RAG7 - Difference
RAG8 - Difference
RAG9 - Difference
RAG10 - Difference
RAG11 - Difference
RAG12 - Difference
RAG13 - Difference
RAG14 - Difference
RAG15 - Difference
RAG16 - Difference
RAG17 - Difference
RAG18 - Difference
RAG19 - Difference
RAG20 - Difference
RAG21 - Difference
RAG22 - Difference
RAG23 - Difference
Horizontal - Vertical
r (mm)
mean $\hat{K}^{iso}_{sector,-15,15}(r) - \hat{K}^{iso}_{sector,75,105}(r)$
$K^{pois}_{sector,-15,15}(r) - K^{pois}_{sector,75,105}(r)$
lo $\hat{K}^{iso}_{sector,-15,15}(r) - \hat{K}^{iso}_{sector,75,105}(r)$
hi $\hat{K}^{iso}_{sector,-15,15}(r) - \hat{K}^{iso}_{sector,75,105}(r)$

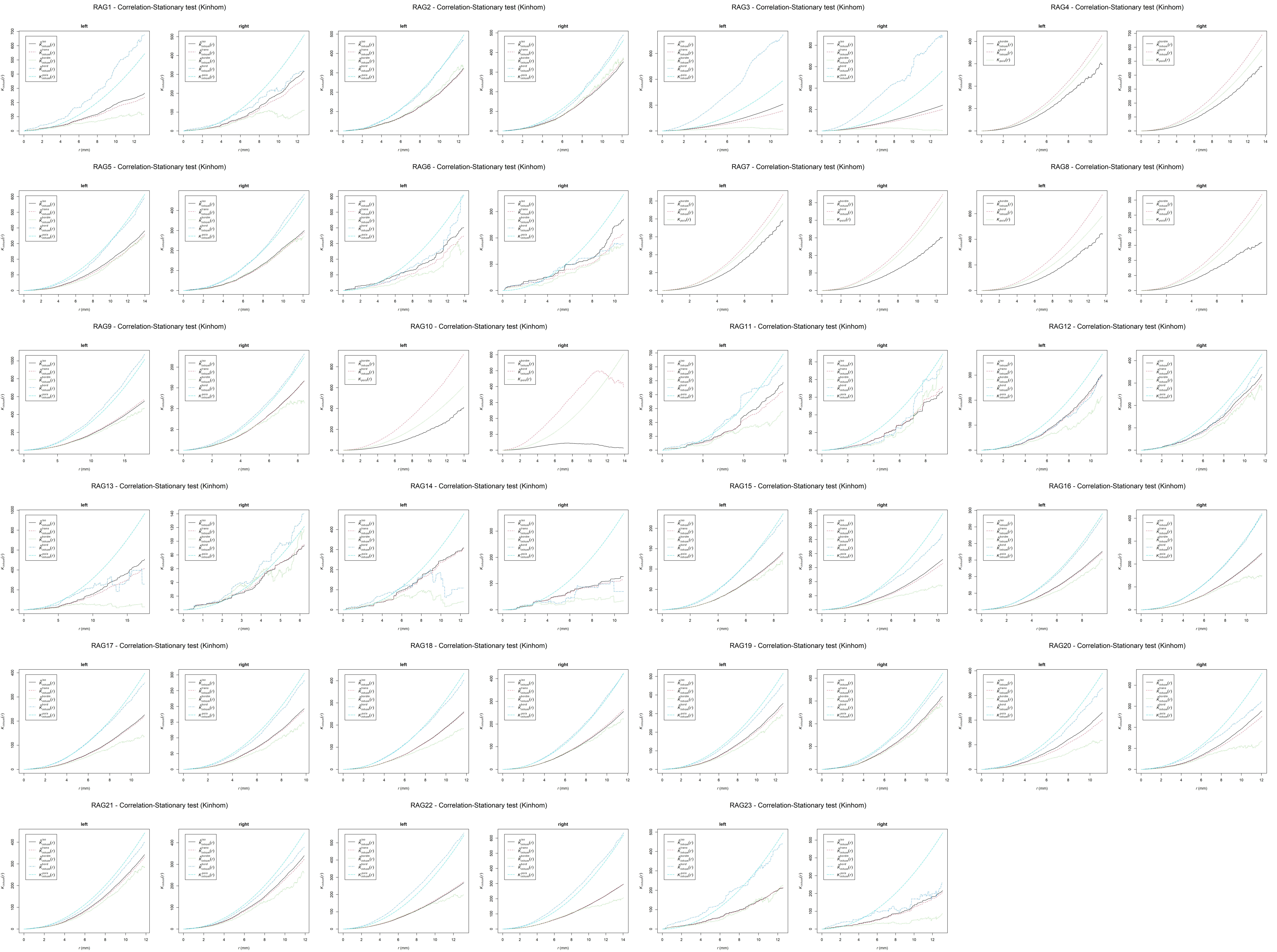
RAG1 - Correlation-Stationary test (Kinhom)
RAG2 - Correlation-Stationary test (Kinhom)
RAG3 - Correlation-Stationary test (Kinhom)
RAG4 - Correlation-Stationary test (Kinhom)
RAG5 - Correlation-Stationary test (Kinhom)
RAG6 - Correlation-Stationary test (Kinhom)
RAG7 - Correlation-Stationary test (Kinhom)
RAG8 - Correlation-Stationary test (Kinhom)
RAG9 - Correlation-Stationary test (Kinhom)
RAG10 - Correlation-Stationary test (Kinhom)
RAG11 - Correlation-Stationary test (Kinhom)
RAG12 - Correlation-Stationary test (Kinhom)
RAG13 - Correlation-Stationary test (Kinhom)
RAG14 - Correlation-Stationary test (Kinhom)
RAG15 - Correlation-Stationary test (Kinhom)
RAG16 - Correlation-Stationary test (Kinhom)
RAG17 - Correlation-Stationary test (Kinhom)
RAG18 - Correlation-Stationary test (Kinhom)
RAG19 - Correlation-Stationary test (Kinhom)
RAG20 - Correlation-Stationary test (Kinhom)
RAG21 - Correlation-Stationary test (Kinhom)
RAG22 - Correlation-Stationary test (Kinhom)
RAG23 - Correlation-Stationary test (Kinhom)
left
right
r (mm)

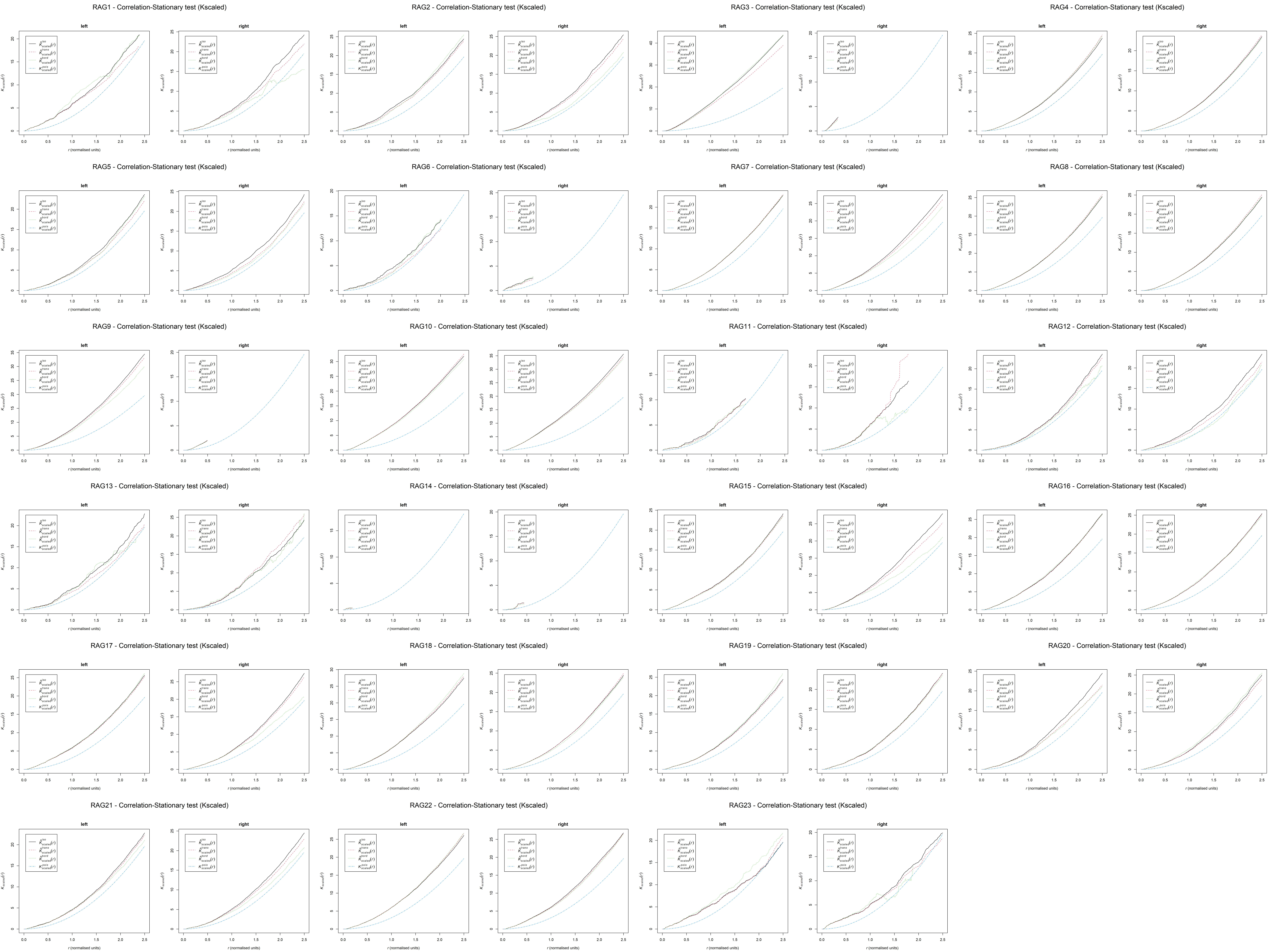
RAG1 - Correlation-Stationary test (Kscaled)
RAG2 - Correlation-Stationary test (Kscaled)
RAG3 - Correlation-Stationary test (Kscaled)
RAG4 - Correlation-Stationary test (Kscaled)
RAG5 - Correlation-Stationary test (Kscaled)
RAG6 - Correlation-Stationary test (Kscaled)
RAG7 - Correlation-Stationary test (Kscaled)
RAG8 - Correlation-Stationary test (Kscaled)
RAG9 - Correlation-Stationary test (Kscaled)
RAG10 - Correlation-Stationary test (Kscaled)
RAG11 - Correlation-Stationary test (Kscaled)
RAG12 - Correlation-Stationary test (Kscaled)
RAG13 - Correlation-Stationary test (Kscaled)
RAG14 - Correlation-Stationary test (Kscaled)
RAG15 - Correlation-Stationary test (Kscaled)
RAG16 - Correlation-Stationary test (Kscaled)
RAG17 - Correlation-Stationary test (Kscaled)
RAG18 - Correlation-Stationary test (Kscaled)
RAG19 - Correlation-Stationary test (Kscaled)
RAG20 - Correlation-Stationary test (Kscaled)
RAG21 - Correlation-Stationary test (Kscaled)
RAG22 - Correlation-Stationary test (Kscaled)
RAG23 - Correlation-Stationary test (Kscaled)
left
right
r (normalised units)

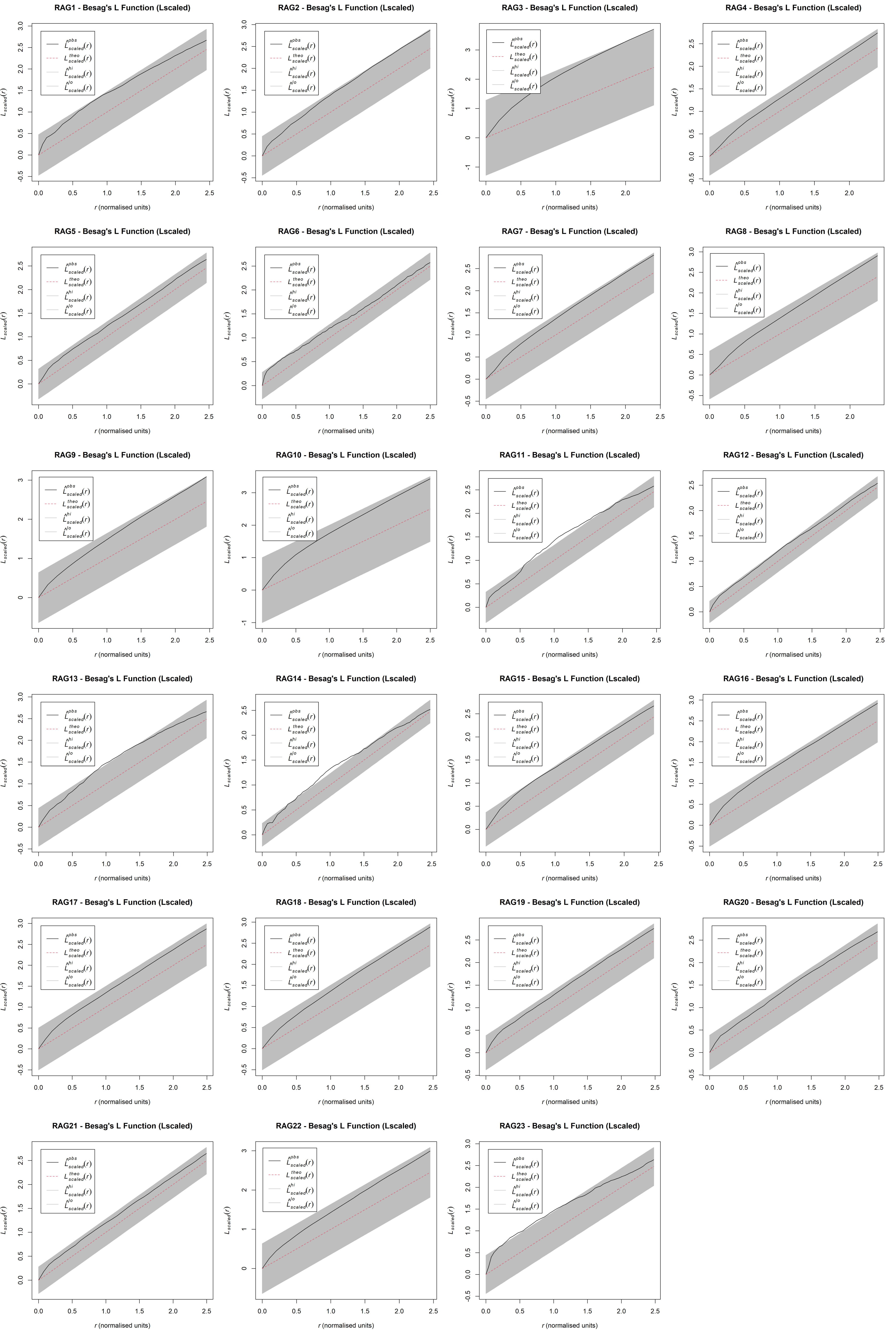
RAG1 - Besag's L Function (Lscaled)
RAG2 - Besag's L Function (Lscaled)
RAG3 - Besag's L Function (Lscaled)
RAG4 - Besag's L Function (Lscaled)
RAG5 - Besag's L Function (Lscaled)
RAG6 - Besag's L Function (Lscaled)
RAG7 - Besag's L Function (Lscaled)
RAG8 - Besag's L Function (Lscaled)
RAG9 - Besag's L Function (Lscaled)
RAG10 - Besag's L Function (Lscaled)
RAG11 - Besag's L Function (Lscaled)
RAG12 - Besag's L Function (Lscaled)
RAG13 - Besag's L Function (Lscaled)
RAG14 - Besag's L Function (Lscaled)
RAG15 - Besag's L Function (Lscaled)
RAG16 - Besag's L Function (Lscaled)
RAG17 - Besag's L Function (Lscaled)
RAG18 - Besag's L Function (Lscaled)
RAG19 - Besag's L Function (Lscaled)
RAG20 - Besag's L Function (Lscaled)
RAG21 - Besag's L Function (Lscaled)
RAG22 - Besag's L Function (Lscaled)
RAG23 - Besag's L Function (Lscaled)
$\hat{L}^{obs}_{scaled}(r)$
$L^{theo}_{scaled}(r)$
$\hat{L}^{hi}_{scaled}(r)$
$\hat{L}^{lo}_{scaled}(r)$
$L_{scaled}(r)$
$r$ (normalised units)

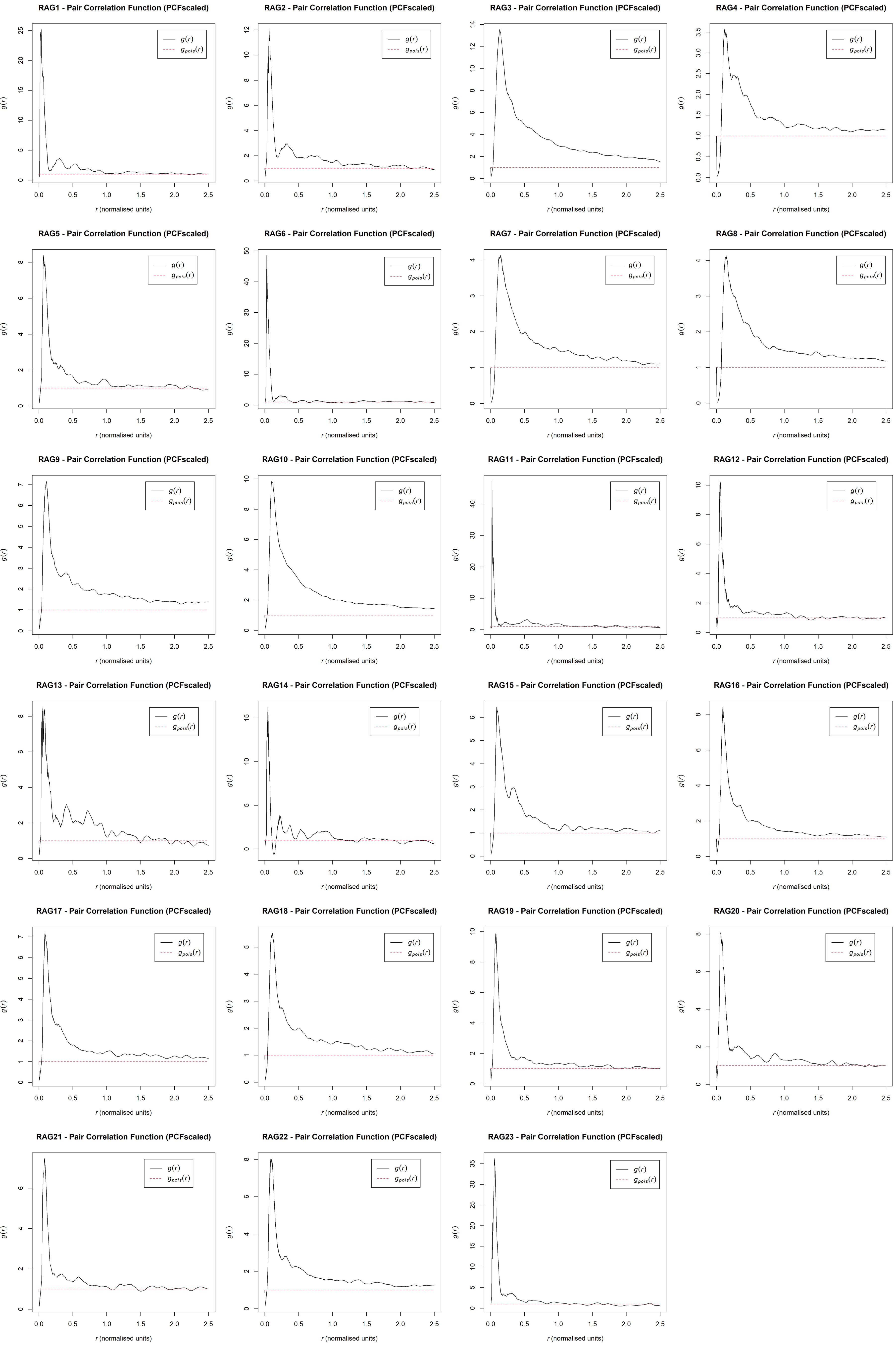
RAG1 - Pair Correlation Function (PCFscaled)
RAG2 - Pair Correlation Function (PCFscaled)
RAG3 - Pair Correlation Function (PCFscaled)
RAG4 - Pair Correlation Function (PCFscaled)
RAG5 - Pair Correlation Function (PCFscaled)
RAG6 - Pair Correlation Function (PCFscaled)
RAG7 - Pair Correlation Function (PCFscaled)
RAG8 - Pair Correlation Function (PCFscaled)
RAG9 - Pair Correlation Function (PCFscaled)
RAG10 - Pair Correlation Function (PCFscaled)
RAG11 - Pair Correlation Function (PCFscaled)
RAG12 - Pair Correlation Function (PCFscaled)
RAG13 - Pair Correlation Function (PCFscaled)
RAG14 - Pair Correlation Function (PCFscaled)
RAG15 - Pair Correlation Function (PCFscaled)
RAG16 - Pair Correlation Function (PCFscaled)
RAG17 - Pair Correlation Function (PCFscaled)
RAG18 - Pair Correlation Function (PCFscaled)
RAG19 - Pair Correlation Function (PCFscaled)
RAG20 - Pair Correlation Function (PCFscaled)
RAG21 - Pair Correlation Function (PCFscaled)
RAG22 - Pair Correlation Function (PCFscaled)
RAG23 - Pair Correlation Function (PCFscaled)
$g(r)$
$g_{pois}(r)$
$r$ (normalised units)

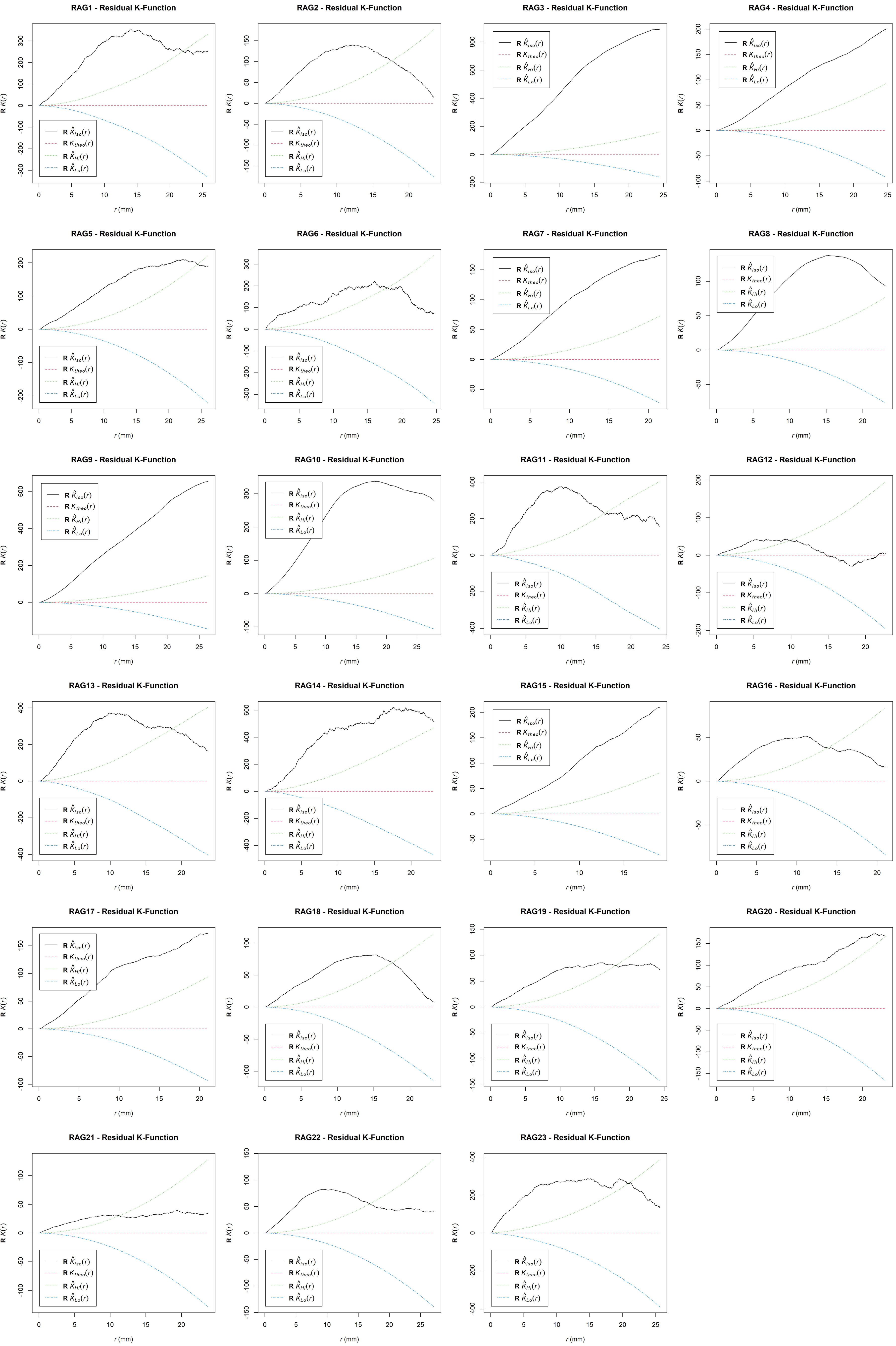
RAG1 - Residual K-Function
RAG2 - Residual K-Function
RAG3 - Residual K-Function
RAG4 - Residual K-Function
RAG5 - Residual K-Function
RAG6 - Residual K-Function
RAG7 - Residual K-Function
RAG8 - Residual K-Function
RAG9 - Residual K-Function
RAG10 - Residual K-Function
RAG11 - Residual K-Function
RAG12 - Residual K-Function
RAG13 - Residual K-Function
RAG14 - Residual K-Function
RAG15 - Residual K-Function
RAG16 - Residual K-Function
RAG17 - Residual K-Function
RAG18 - Residual K-Function
RAG19 - Residual K-Function
RAG20 - Residual K-Function
RAG21 - Residual K-Function
RAG22 - Residual K-Function
RAG23 - Residual K-Function
R $K(r)$
$r$ (mm)
R $\hat{K}_{iso}(r)$
R $K_{theo}(r)$
R $\hat{K}_{Hi}(r)$
R $\hat{K}_{Lo}(r)$

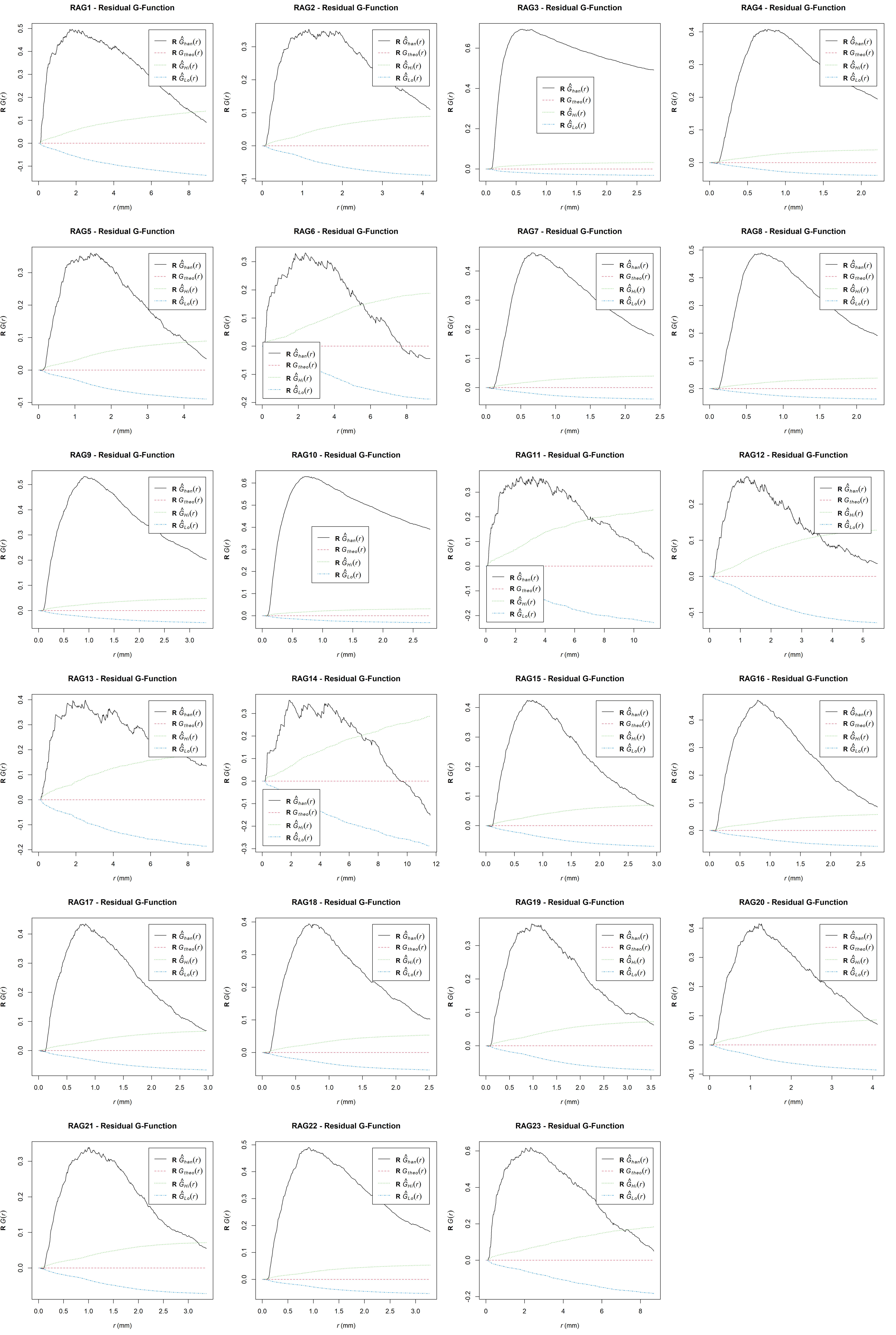

RAG1 - Residual G-Function
RAG2 - Residual G-Function
RAG3 - Residual G-Function
RAG4 - Residual G-Function
RAG5 - Residual G-Function
RAG6 - Residual G-Function
RAG7 - Residual G-Function
RAG8 - Residual G-Function
RAG9 - Residual G-Function
RAG10 - Residual G-Function
RAG11 - Residual G-Function
RAG12 - Residual G-Function
RAG13 - Residual G-Function
RAG14 - Residual G-Function
RAG15 - Residual G-Function
RAG16 - Residual G-Function
RAG17 - Residual G-Function
RAG18 - Residual G-Function
RAG19 - Residual G-Function
RAG20 - Residual G-Function
RAG21 - Residual G-Function
RAG22 - Residual G-Function
RAG23 - Residual G-Function
R $G(r)$
$r$ (mm)
R $\hat{G}_{han}(r)$
R $G_{theo}(r)$
R $\hat{G}_{Hi}(r)$
R $\hat{G}_{Lo}(r)$

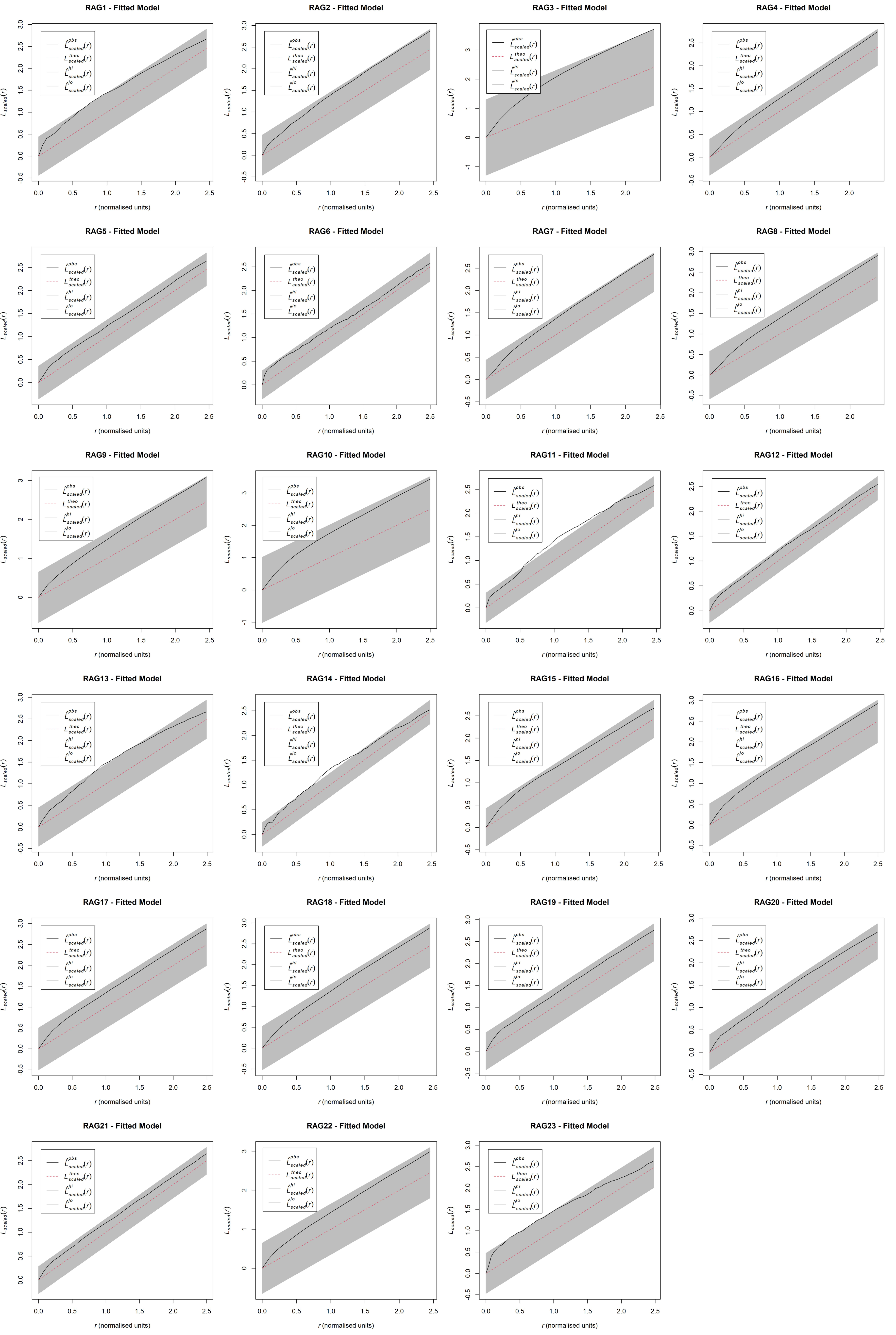
RAG1 - Fitted Model
RAG2 - Fitted Model
RAG3 - Fitted Model
RAG4 - Fitted Model
RAG5 - Fitted Model
RAG6 - Fitted Model
RAG7 - Fitted Model
RAG8 - Fitted Model
RAG9 - Fitted Model
RAG10 - Fitted Model
RAG11 - Fitted Model
RAG12 - Fitted Model
RAG13 - Fitted Model
RAG14 - Fitted Model
RAG15 - Fitted Model
RAG16 - Fitted Model
RAG17 - Fitted Model
RAG18 - Fitted Model
RAG19 - Fitted Model
RAG20 - Fitted Model
RAG21 - Fitted Model
RAG22 - Fitted Model
RAG23 - Fitted Model
$\hat{L}^{obs}_{scaled}(r)$
$L^{theo}_{scaled}(r)$
$\hat{L}^{hi}_{scaled}(r)$
$\hat{L}^{lo}_{scaled}(r)$
$L_{scaled}(r)$
$r$ (normalised units)

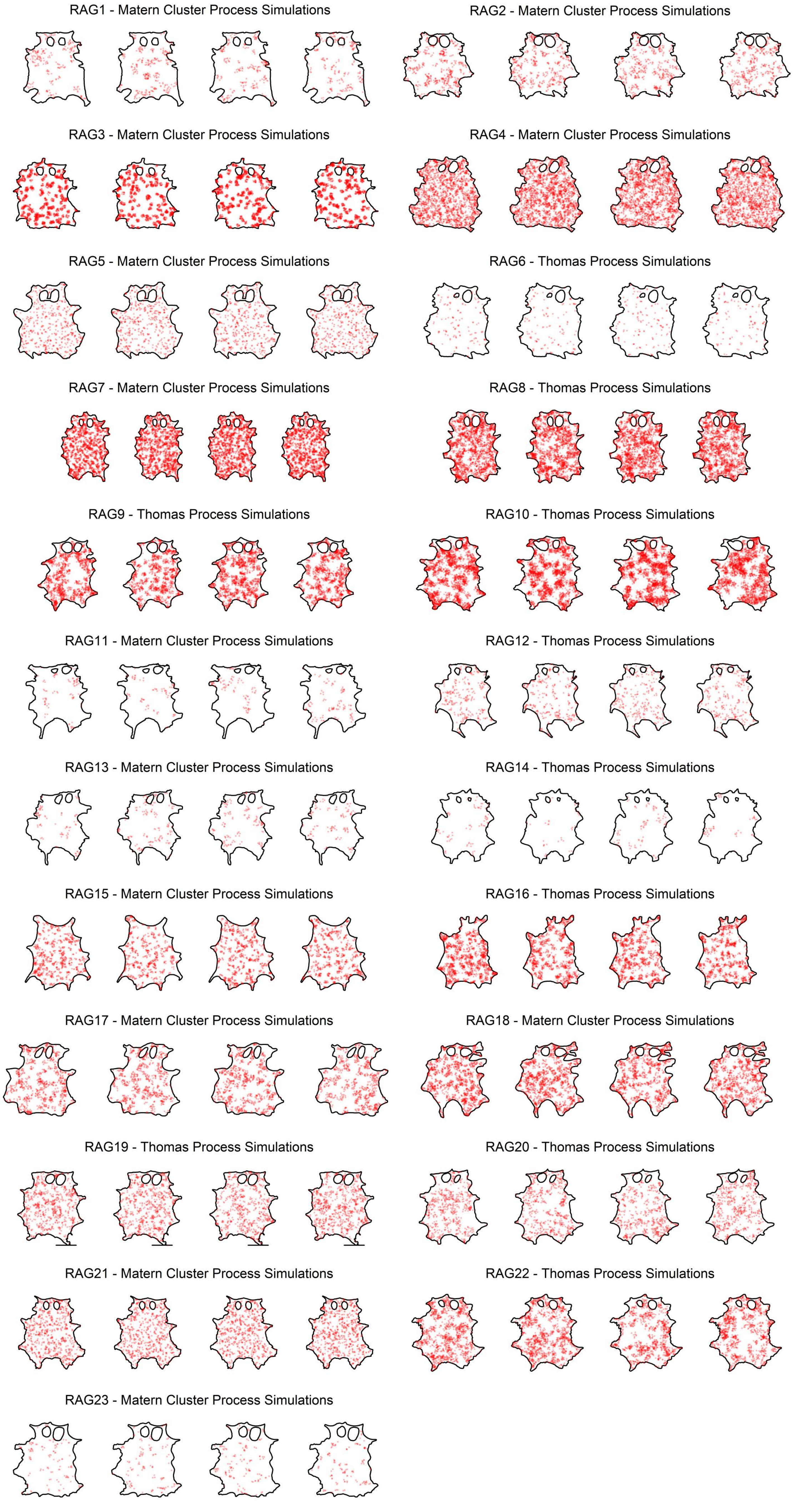
RAG1 - Matern Cluster Process Simulations
RAG2 - Matern Cluster Process Simulations
RAG3 - Matern Cluster Process Simulations
RAG4 - Matern Cluster Process Simulations
RAG5 - Matern Cluster Process Simulations
RAG6 - Thomas Process Simulations
RAG7 - Matern Cluster Process Simulations
RAG8 - Thomas Process Simulations
RAG9 - Thomas Process Simulations
RAG10 - Thomas Process Simulations
RAG11 - Matern Cluster Process Simulations
RAG12 - Thomas Process Simulations
RAG13 - Matern Cluster Process Simulations
RAG14 - Thomas Process Simulations
RAG15 - Matern Cluster Process Simulations
RAG16 - Thomas Process Simulations
RAG17 - Matern Cluster Process Simulations
RAG18 - Matern Cluster Process Simulations
RAG19 - Thomas Process Simulations
RAG20 - Thomas Process Simulations
RAG21 - Matern Cluster Process Simulations
RAG22 - Thomas Process Simulations
RAG23 - Matern Cluster Process Simulations

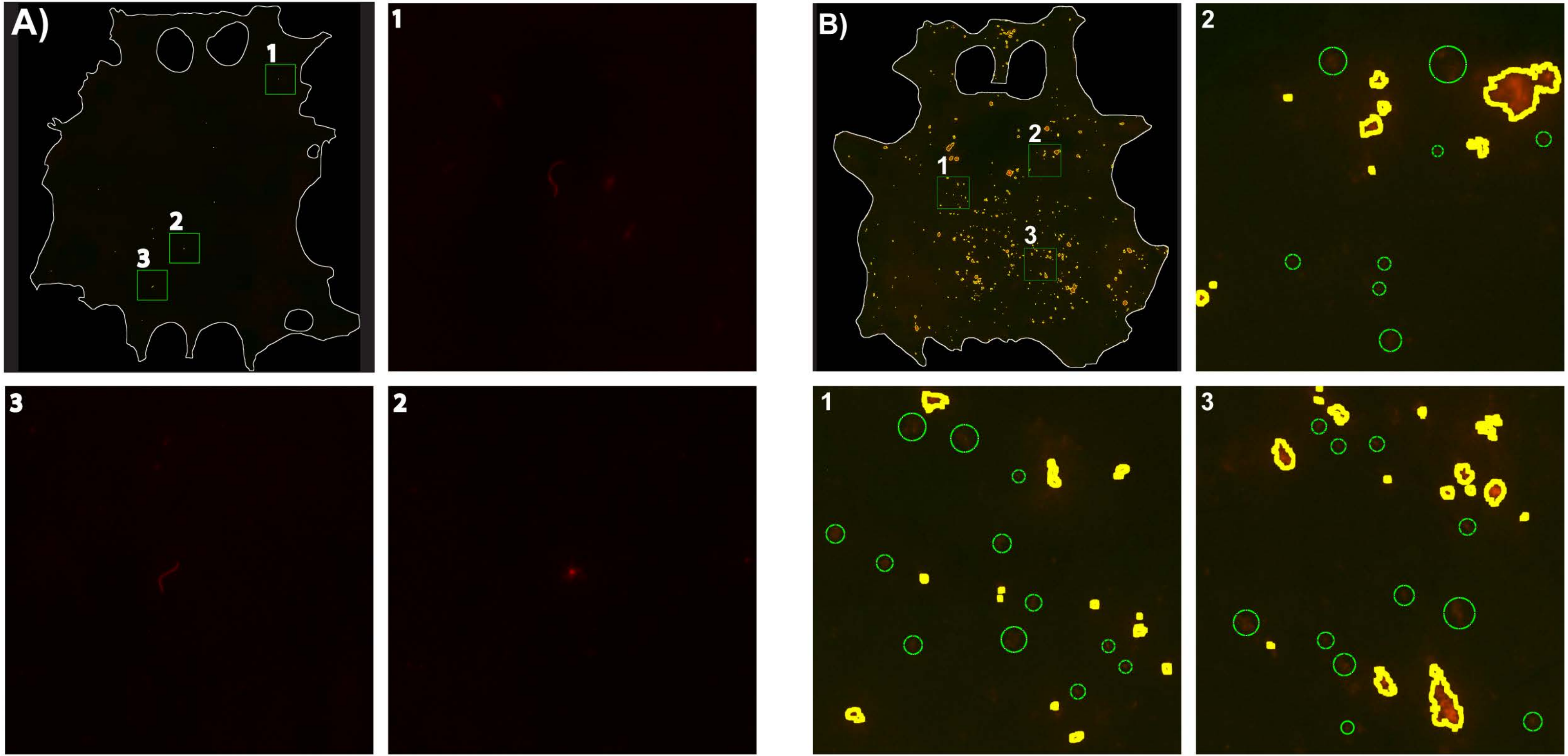
A)
1
2
3
B)
1
2
3

### Table S1

*Statistical information on the skin parasite patch analysis*

| | Patch Areas | | | | | | | | | | | | | | | | | |
|---|---|---|---|---|---|---|---|---|---|---|---|---|---|---|---|---|---|---|
| Mouse IDs | Min. (μm$^2$) | 1$^{st}$ Quarter (μm$^2$) | Median (μm$^2$) | Mean (μm$^2$) | 3$^{rd}$ Quarter (μm$^2$) | Max. (μm$^2$) | Linear, P-Value | Outcome | Log, P-Value | Outcome | Skewness | Outcome | S.E. of Skewness | Test Statistic | Outcome | Total Patch Area (cm$^2$) | Total Skin Area (cm$^2$) | Patch Ratio of Skin (%) |
| **RAG1** | 4150.4 | 6456.1 | 12912.2 | 133496.4 | 53032.4 | 6942633.5 | <0.0001 | Non-Normal | <0.0001 | Non-Normal | 10.13 | Skewed | 0.159 | 63.799 | Skewed | 0.314 | 78.212 | 0.401 |
| **RAG2** | 4150.4 | 5995.0 | 11528.8 | 34514.7 | 27207.9 | 2634557.6 | <0.0001 | Non-Normal | <0.0001 | Non-Normal | 14.83 | Skewed | 0.111 | 133.469 | Skewed | 0.167 | 57.729 | 0.289 |
| **RAG3** | 4150.4 | 5072.7 | 6917.3 | 3027315.9 | 13373.4 | 5310213076.6 | <0.0001 | Non-Normal | <0.0001 | Non-Normal | 42.25 | Skewed | 0.058 | 730.571 | Skewed | 54.219 | 66.828 | 81.132 |
| **RAG4** | 4189.1 | 7214.6 | 15942.0 | 534472.1 | 61178.8 | 286399016.2 | <0.0001 | Non-Normal | <0.0001 | Non-Normal | 28.16 | Skewed | 0.053 | 533.432 | Skewed | 11.491 | 69.068 | 16.637 |
| **RAG5** | 4189.1 | 9454.6 | 21003.9 | 111904.8 | 84190.0 | 2603663.4 | <0.0001 | Non-Normal | <0.0001 | Non-Normal | 5.51 | Skewed | 0.109 | 50.750 | Skewed | 0.566 | 73.492 | 0.770 |
| **RAG6** | 4189.1 | 6749.2 | 13032.9 | 38610.9 | 31069.4 | 586478.8 | <0.0001 | Non-Normal | <0.0001 | Non-Normal | 4.81 | Skewed | 0.212 | 22.730 | Skewed | 0.051 | 77.209 | 0.066 |
| **RAG7** | 4189.1 | 6632.8 | 13382.0 | 290149.8 | 47476.9 | 173631707.6 | <0.0001 | Non-Normal | <0.0001 | Non-Normal | 37.49 | Skewed | 0.053 | 711.487 | Skewed | 6.261 | 70.762 | 8.849 |
| **RAG8** | 4189.1 | 6981.9 | 13498.3 | 383602.7 | 42473.2 | 314545112.0 | <0.0001 | Non-Normal | <0.0001 | Non-Normal | 35.30 | Skewed | 0.051 | 698.310 | Skewed | 8.995 | 71.296 | 12.617 |
| **RAG9** | 4189.1 | 6167.3 | 12683.8 | 254524.9 | 44335.0 | 86460824.7 | <0.0001 | Non-Normal | <0.0001 | Non-Normal | 24.02 | Skewed | 0.064 | 374.435 | Skewed | 3.703 | 80.021 | 4.628 |
| **RAG10** | 4189.1 | 5469.1 | 8611.0 | 1882972.3 | 20480.2 | 4361937113.3 | <0.0001 | Non-Normal | <0.0001 | Non-Normal | 49.50 | Skewed | 0.049 | 1002.298 | Skewed | 46.265 | 82.409 | 56.140 |
| **RAG11** | 4305.5 | 8320.1 | 20596.6 | 214709.3 | 72029.8 | 10471439.6 | <0.0001 | Non-Normal | <0.0001 | Non-Normal | 8.87 | Skewed | 0.233 | 38.148 | Skewed | 0.232 | 62.160 | 0.373 |
| **RAG12** | 4189.1 | 8000.1 | 18967.5 | 62696.1 | 49455.1 | 5809980.3 | <0.0001 | Non-Normal | <0.0001 | Non-Normal | 15.81 | Skewed | 0.146 | 108.579 | Skewed | 0.176 | 57.105 | 0.307 |
| **RAG13** | 4189.1 | 9949.2 | 30720.3 | 159295.1 | 84655.4 | 3995619.6 | <0.0001 | Non-Normal | <0.0001 | Non-Normal | 5.65 | Skewed | 0.206 | 27.485 | Skewed | 0.221 | 58.000 | 0.382 |
| **RAG14** | 4189.1 | 9541.9 | 21527.5 | 49925.1 | 49193.2 | 705287.3 | <0.0001 | Non-Normal | 0.0266 | Non-Normal | 5.25 | Skewed | 0.276 | 19.047 | Skewed | 0.038 | 64.535 | 0.059 |
| **RAG15** | 4189.1 | 6138.2 | 11229.2 | 433597.0 | 58589.7 | 70715614.0 | <0.0001 | Non-Normal | <0.0001 | Non-Normal | 14.55 | Skewed | 0.086 | 168.742 | Skewed | 3.486 | 40.401 | 8.629 |
| **RAG16** | 4189.1 | 6167.3 | 11403.8 | 256916.4 | 43520.4 | 23751228.3 | <0.0001 | Non-Normal | <0.0001 | Non-Normal | 10.22 | Skewed | 0.072 | 141.489 | Skewed | 2.947 | 51.233 | 5.752 |
| **RAG17** | 4189.1 | 6400.1 | 12800.1 | 232102.4 | 64524.3 | 20451586.8 | <0.0001 | Non-Normal | <0.0001 | Non-Normal | 10.17 | Skewed | 0.082 | 123.584 | Skewed | 2.049 | 47.587 | 4.307 |
| **RAG18** | 4189.1 | 6749.2 | 14312.9 | 252552.2 | 55506.0 | 53973854.8 | <0.0001 | Non-Normal | <0.0001 | Non-Normal | 19.34 | Skewed | 0.068 | 285.988 | Skewed | 3.306 | 56.286 | 5.873 |
| **RAG19** | 4189.1 | 6981.9 | 20014.8 | 741849.7 | 179027.3 | 91466142.1 | <0.0001 | Non-Normal | <0.0001 | Non-Normal | 14.92 | Skewed | 0.090 | 166.476 | Skewed | 5.519 | 64.264 | 8.589 |
| **RAG20** | 4189.1 | 8843.7 | 33280.3 | 481798.2 | 210678.5 | 29861546.3 | <0.0001 | Non-Normal | <0.0001 | Non-Normal | 10.27 | Skewed | 0.105 | 97.610 | Skewed | 2.597 | 56.540 | 4.593 |
| **RAG21** | 4189.1 | 8611.0 | 28218.5 | 658429.5 | 128030.4 | 113190876.7 | <0.0001 | Non-Normal | <0.0001 | Non-Normal | 17.22 | Skewed | 0.088 | 196.464 | Skewed | 5.123 | 59.912 | 8.550 |
| **RAG22** | 4150.4 | 6917.3 | 17984.9 | 1410841.9 | 75167.7 | 1053270594.4 | <0.0001 | Non-Normal | <0.0001 | Non-Normal | 31.89 | Skewed | 0.069 | 459.555 | Skewed | 17.537 | 73.533 | 23.849 |
| **RAG23** | 4150.4 | 8300.7 | 21674.1 | 111772.8 | 103759.1 | 1681819.0 | <0.0001 | Non-Normal | <0.0001 | Non-Normal | 4.34 | Skewed | 0.203 | 21.408 | Skewed | 0.160 | 72.675 | 0.220 |

| | Skin Parasites | | | | | | | | | | | | | | Patch Densities | | | | | Patch Counts | |
|---|---|---|---|---|---|---|---|---|---|---|---|---|---|---|---|---|---|---|---|---|---|
| Mouse IDs | Min. | 1$^{st}$ Quarter | Median | Mean | 3$^{rd}$ Quarter | Max. | Linear, P-Value | Outcome | Log, P-Value | Outcome | per cm$^2$ | per Skin (cm$^2$) | | | Skewness | Outcome | S.E. of Skewness | Test Statistic | Outcome | per cm$^2$ | per skin |
| **RAG1** | 251.0 | 2290.0 | 6648.1 | 10570.2 | 12629.7 | 43123.5 | 0.0001 | Non-Normal | 0.6208 | Normal | 23513.0 | 1839000 | | | 2.35 | Skewed | 0.159 | 14.800 | Skewed | 3.0 | 235 |
| **RAG2** | 1358.6 | 3503.0 | 5528.4 | 9424.0 | 9158.3 | 79325.4 | <0.0001 | Non-Normal | 0.1697 | Normal | 19552.7 | 1128754 | | | 1.25 | Skewed | 0.111 | 11.250 | Skewed | 8.4 | 483 |
| **RAG3** | 193055.3 | 3662059.8 | 5266551.5 | 8605329.5 | 9368950.5 | 34302760.0 | 0.0002 | Non-Normal | 0.0945 | Normal | 18626615.7 | 1244782320 | | | 8.65 | Skewed | 0.058 | 149.572 | Skewed | 26.8 | 1791 |
| **RAG4** | 6164.2 | 23319.2 | 39409.3 | 67356.2 | 99910.3 | 335843.0 | <0.0001 | Non-Normal | 0.8846 | Normal | 139381.9 | 9626886 | | | 2.16 | Skewed | 0.053 | 40.917 | Skewed | 31.1 | 2150 |
| **RAG5** | 630.3 | 2643.5 | 4830.5 | 21383.9 | 11606.3 | 287609.7 | <0.0001 | Non-Normal | 0.1611 | Normal | 17084.4 | 1255562 | | | 1.64 | Skewed | 0.109 | 15.105 | Skewed | 6.9 | 506 |
| **RAG6** | 2780.3 | 5588.8 | 9640.3 | 16032.1 | 15559.0 | 82828.4 | <0.0001 | Non-Normal | 0.4119 | Normal | 34095.8 | 2632497 | | | 3.42 | Skewed | 0.212 | 16.161 | Skewed | 1.7 | 131 |
| **RAG7** | 6126.3 | 49168.5 | 78845.5 | 119731.5 | 163596.8 | 353988.8 | 0.0014 | Non-Normal | 0.1613 | Normal | 278859.1 | 19732605 | | | 2.34 | Skewed | 0.053 | 44.409 | Skewed | 30.5 | 2158 |
| **RAG8** | 3933.0 | 41787.6 | 77590.1 | 93149.6 | 120658.0 | 247062.3 | 0.2359 | Normal | 0.0256 | Non-Normal | 274418.8 | 19564890 | | | 2.73 | Skewed | 0.051 | 54.005 | Skewed | 32.9 | 2345 |
| **RAG9** | 473.9 | 11507.2 | 26347.2 | 45588.5 | 48604.2 | 181003.6 | 0.0012 | Non-Normal | 0.3487 | Normal | 93184.3 | 7456728 | | | 3.82 | Skewed | 0.064 | 59.548 | Skewed | 18.2 | 1455 |
| **RAG10** | 17819.6 | 189430.1 | 469938.6 | 745577.4 | 846442.4 | 2548114.0 | 0.0019 | Non-Normal | 0.6373 | Normal | 1662067.6 | 136969139 | | | 6.07 | Skewed | 0.049 | 122.908 | Skewed | 29.8 | 2457 |
| **RAG11** | 982.4 | 3056.5 | 5428.4 | 12199.7 | 11178.5 | 69469.2 | <0.0001 | Non-Normal | 0.8334 | Normal | 19199.2 | 1193415 | | | 1.87 | Skewed | 0.233 | 8.042 | Skewed | 1.7 | 108 |
| **RAG12** | 1309.1 | 3249.3 | 5024.9 | 5847.2 | 7925.7 | 15235.3 | 0.0943 | Normal | 0.9993 | Normal | 17771.9 | 1014865 | | | 2.83 | Skewed | 0.146 | 19.436 | Skewed | 4.9 | 280 |
| **RAG13** | 2056.1 | 4884.9 | 11918.5 | 13116.8 | 20004.5 | 34186.7 | 0.1923 | Normal | 0.1010 | Normal | 42152.9 | 2444884 | | | 1.79 | Skewed | 0.206 | 8.708 | Skewed | 2.4 | 139 |
| **RAG14** | 1433.8 | 3116.7 | 4589.6 | 5488.3 | 7350.7 | 13219.8 | 0.0299 | Non-Normal | 0.8777 | Normal | 16232.6 | 1047571 | | | 2.11 | Skewed | 0.276 | 7.655 | Skewed | 1.2 | 76 |
| **RAG15** | 4131.5 | 8386.6 | 17612.0 | 19527.3 | 30196.7 | 35784.0 | 0.0367 | Non-Normal | 0.0676 | Normal | 62289.9 | 2516581 | | | 2.62 | Skewed | 0.086 | 30.385 | Skewed | 19.9 | 804 |
| **RAG16** | 7056.4 | 15003.7 | 37690.8 | 38286.2 | 52103.9 | 96301.9 | 0.1778 | Normal | 0.3773 | Normal | 133304.1 | 6829598 | | | 2.77 | Skewed | 0.072 | 38.349 | Skewed | 22.4 | 1147 |
| **RAG17** | 3428.2 | 5491.5 | 9724.7 | 12697.1 | 14388.8 | 44514.5 | 0.0014 | Non-Normal | 0.7918 | Normal | 34393.9 | 1636704 | | | 2.41 | Skewed | 0.082 | 29.286 | Skewed | 18.6 | 883 |
| **RAG18** | 0.0 | 5010.2 | 22375.0 | 26836.9 | 41955.1 | 70036.3 | 0.0752 | Normal | 0.0001 | Non-Normal | 79135.2 | 4454194 | | | 2.64 | Skewed | 0.068 | 39.039 | Skewed | 23.3 | 1309 |
| **RAG19** | 66263.1 | 182743.0 | 311323.8 | 325422.6 | 378153.2 | 802803.1 | 0.0574 | Normal | 0.6473 | Normal | 1101082.7 | 70760337 | | | 2.21 | Skewed | 0.090 | 24.659 | Skewed | 11.6 | 744 |
| **RAG20** | 35844.2 | 144539.6 | 232849.0 | 305971.0 | 337057.3 | 1375823.9 | 0.0001 | Non-Normal | 0.8655 | Normal | 823534.7 | 46562927 | | | 2.55 | Skewed | 0.105 | 24.236 | Skewed | 9.5 | 539 |
| **RAG21** | 49846.0 | 158783.8 | 353309.4 | 747667.7 | 524852.8 | 5646642.5 | <0.0001 | Non-Normal | 0.4689 | Normal | 1249576.4 | 74865201 | | | 2.20 | Skewed | 0.088 | 25.100 | Skewed | 13.0 | 778 |
| **RAG22** | 12411.1 | 319686.2 | 432618.4 | 1011280.0 | 718135.4 | 5146964.0 | <0.0001 | Non-Normal | 0.0947 | Normal | 1530074.8 | 112511713 | | | 3.45 | Skewed | 0.069 | 49.717 | Skewed | 16.9 | 1243 |
| **RAG23** | 23671.2 | 64577.8 | 84107.0 | 86370.6 | 93306.0 | 232578.5 | 0.0126 | Non-Normal | 0.3278 | Normal | 297467.8 | 21618395 | | | 3.82 | Skewed | 0.203 | 18.843 | Skewed | 2.0 | 143 |

### Table S2

*Collective information on the point pattern analysis of skin parasite patch patterns*

| | | Point Pattern Intensity | | | | Quadrat Test of Inhomogeneity | | | | Poisson Process Test | | |
|---|---|---|---|---|---|---|---|---|---|---|---|---|
| Mouse IDs | Double Point Check | Intensity w/o Mark (/ $cm^2$) | S.E. w/o Mark | Intensity by Area (/ $cm^2$) | S.E. by Area | Chi-Square P-values | Fisher's Exact P-values | Accepted Test | Outcome | P-values | Acceted Test | Outcome |
| **RAG1** | No Doubles | 3.014 | 1.168 | 0.404 | 0.324 | 0.002 | 0.001 | Fisher's Exact | Inhomogeneous | <0.0001 | Quadrat Count | Not Poisson |
| **RAG2** | No Doubles | 8.265 | 2.655 | 0.270 | 0.412 | 0.002 | 0.001 | Fisher's Exact | Inhomogeneous | <0.0001 | Quadrat Count | Not Poisson |
| **RAG3** | No Doubles | 26.173 | 9.549 | 80.397 | 3.197 | 0.002 | 0.001 | Fisher's Exact | Inhomogeneous | <0.0001 | Quadrat Count | Not Poisson |
| **RAG4** | No Doubles | 31.007 | 10.572 | 16.624 | 4.526 | 0.002 | 0.001 | Fisher's Exact | Inhomogeneous | <0.0001 | Quadrat Count | Not Poisson |
| **RAG5** | No Doubles | 6.761 | 2.821 | 0.776 | 0.806 | 0.002 | 0.001 | Fisher's Exact | Inhomogeneous | <0.0001 | Quadrat Count | Not Poisson |
| **RAG6** | No Doubles | 1.628 | 0.636 | 0.064 | 0.114 | 0.004 | 0.001 | Fisher's Exact | Inhomogeneous | <0.0001 | Quadrat Count | Not Poisson |
| **RAG7** | No Doubles | 30.374 | 10.016 | 8.892 | 3.605 | 0.002 | 0.001 | Fisher's Exact | Inhomogeneous | <0.0001 | Quadrat Count | Not Poisson |
| **RAG8** | No Doubles | 32.743 | 13.619 | 12.595 | 4.261 | 0.002 | 0.001 | Fisher's Exact | Inhomogeneous | <0.0001 | Quadrat Count | Not Poisson |
| **RAG9** | No Doubles | 18.081 | 5.212 | 4.587 | 1.735 | 0.002 | 0.001 | Fisher's Exact | Inhomogeneous | <0.0001 | Quadrat Count | Not Poisson |
| **RAG10** | No Doubles | 29.583 | 14.797 | 56.147 | 4.086 | 0.002 | 0.001 | Fisher's Exact | Inhomogeneous | <0.0001 | Quadrat Count | Not Poisson |
| **RAG11** | No Doubles | 1.734 | 0.661 | 0.329 | 0.204 | 0.002 | 0.001 | Fisher's Exact | Inhomogeneous | <0.0001 | Quadrat Count | Not Poisson |
| **RAG12** | No Doubles | 4.896 | 1.794 | 0.262 | 0.359 | 0.008 | 0.001 | Fisher's Exact | Inhomogeneous | <0.0001 | Quadrat Count | Not Poisson |
| **RAG13** | No Doubles | 2.414 | 0.995 | 0.396 | 0.315 | 0.002 | 0.001 | Fisher's Exact | Inhomogeneous | <0.0001 | Quadrat Count | Not Poisson |
| **RAG14** | No Doubles | 1.188 | 0.450 | 0.059 | 0.089 | 0.002 | 0.001 | Fisher's Exact | Inhomogeneous | <0.0001 | Quadrat Count | Not Poisson |
| **RAG15** | No Doubles | 19.904 | 8.774 | 8.931 | 3.396 | 0.002 | 0.001 | Fisher's Exact | Inhomogeneous | <0.0001 | Quadrat Count | Not Poisson |
| **RAG16** | No Doubles | 22.377 | 8.678 | 5.820 | 3.097 | 0.002 | 0.001 | Fisher's Exact | Inhomogeneous | <0.0001 | Quadrat Count | Not Poisson |
| **RAG17** | No Doubles | 18.576 | 7.716 | 4.286 | 2.707 | 0.002 | 0.001 | Fisher's Exact | Inhomogeneous | <0.0001 | Quadrat Count | Not Poisson |
| **RAG18** | No Doubles | 23.285 | 9.008 | 5.951 | 3.114 | 0.002 | 0.001 | Fisher's Exact | Inhomogeneous | <0.0001 | Quadrat Count | Not Poisson |
| **RAG19** | No Doubles | 11.549 | 4.256 | 8.654 | 2.746 | 0.002 | 0.001 | Fisher's Exact | Inhomogeneous | <0.0001 | Quadrat Count | Not Poisson |
| **RAG20** | No Doubles | 9.473 | 4.018 | 4.627 | 2.132 | 0.002 | 0.001 | Fisher's Exact | Inhomogeneous | <0.0001 | Quadrat Count | Not Poisson |
| **RAG21** | No Doubles | 13.021 | 4.663 | 8.812 | 2.727 | 0.002 | 0.001 | Fisher's Exact | Inhomogeneous | <0.0001 | Quadrat Count | Not Poisson |
| **RAG22** | No Doubles | 16.866 | 7.165 | 23.822 | 3.450 | 0.002 | 0.001 | Fisher's Exact | Inhomogeneous | <0.0001 | Quadrat Count | Not Poisson |
| **RAG23** | No Doubles | 1.948 | 0.880 | 0.215 | 0.246 | 0.002 | 0.001 | Fisher's Exact | Inhomogeneous | <0.0001 | Quadrat Count | Not Poisson |

### Table S3

*Model parameter ranges*

| | kappa | | scale | |
|---|---|---|---|---|
| **Models** | **Min kappa** | **Max kappa** | **Min scale** | **Max scale** |
| **Log-Gaussian Cox Process** | 0.0005264 | 0.0790058 | 0.3797 | 2.7016 |
| **Cauchy Cluster Process** | 0.7338 | 4.8284 | 1.160 | 18.991 |
| **Matern Cluster Process** | 0.002702 | 0.105462 | 1.126 | 5.768 |
| **Thomas Cluster Process** | 0.002256 | 0.128535 | 0.5816 | 2.6680 |
| **Variance Gamma Cluster Process** | 0.0006041 | 0.0955190 | 0.649 | 4.063 |
